# Genotype-specific nonphotochemical quenching responses to nitrogen deficit are linked to chlorophyll a to b ratios

**DOI:** 10.1101/2023.10.24.563650

**Authors:** Seema Sahay, Marcin Grzybowski, James C. Schnable, Katarzyna Głowacka

## Abstract

Non-photochemical quenching (NPQ) protects plants from photodamage caused by excess light energy. The mechanism of NPQ appears to be conserved across flowering plants. However, substantial variation in NPQ has been reported within different genotypes of the same species grown under the environmental conditions. Individual maize genotypes from a diversity panel exhibited a range of responses to low nitrogen with some genotypes exhibiting increased NPQ between control and low nitrogen conditions and others exhibiting no change. These patterns were consistent for the same genotypes across multiple field seasons. NPQ increases under low nitrogen were correlated with shifts in the ratio of chlorophyll *a* to chlorophyll *b* consistent with a decrease in reaction centers. Both photosynthetic capacity and dry biomass accumulation decreased more in maize genotypes which were unable to maintain constant NPQ levels between control and low N conditions. Collectively these results suggest that the ability to maintain sufficient numbers of reaction centers under low nitrogen conditions and avoid dissipating a greater proportion of absorbed light energy via the NPQ pathway may play a key role in increasing carbon fixation and productivity in nitrogen-limited environments.

**Highlights:**

- Substantial variation in NPQ kinetics exists in maize on both low and control N.
- In early and late-vegetative stages a similar portion of genotypes increased, no-change or decreased *NPQ_max_* in response to low N while in the post-flowering stage substantially more genotypes decreased *NPQ_max_*.
- In low nitrogen conditions, the *NPQ_max_* strongly correlates with shifts in Chl *a*/Chl *b* ratios.

## Introduction

The production and application of synthetic nitrogen (N) fertilizer have played a critical role in our civilization’s ability to meet growing needs for food and fuel over the last century. Prior to the advent of synthetically produced biologically accessible forms of nitrogen, access to nitrogen was the rate-limiting factor on crop productivity in many environments. Today, the use of more than 118 million tons of N fertilizer per year by the agricultural sector (FAO 2019; Sandhu et al., 2021) is responsible for half the calories used to feed the world’s population (Smil, 2001). However, the production of this fertilizer is energy intensive (Giles, 2005; Hirel *et al*., 2007) and the application of N fertilizer to fields results in the production of nitrous oxide (N_2_O) a greenhouse gas estimated to be 298 times more potent than CO_2_ (Stulen *et al*., 1998; Baggs *et al*., 2006; Park *et al*., 2012). One of the crops which has benefited substantially from access to N fertilizer is maize (Duvick, 2005; USDA-NASS, 2017). Today many farmers apply nitrogen fertilizer to maize fields in excess of 300 kg/ha/year (Zhang *et al*., 2015; Cao *et al*., 2018). The cost to farmers of nitrogen fertilizer can exceed $300 ha/year (‘Statista 2022’; Rodríguez-Espinosa *et al*., 2023).

Given both the environmental and economic costs of excessive use of nitrogen, there is growing interest in breeding maize varieties that are more productive under nitrogen limited conditions. However, there are many biological challenges which must be overcome to achieve this goal (Ciampitti *et al*., 2022). One of these challenges is that nitrogen deficit presents substantial challenges to maintaining plant photosynthetic productivity. Low N availability hinders the growth and development of plants at both physiological and molecular levels (Martin *et al*., 2002; Ding *et al*., 2005; Peng *et al*., 2007; Kant *et al*., 2008; Bi *et al*., 2009). Nitrogen deficiency accelerates leaf yellowing via chlorophyll degradation, anthocyanin accumulation and senescence which decreases the rate of CO_2_ assimilation and reduces soluble protein content (Ding *et al*., 2005; Diaz *et al*., 2006; Zhao *et al*., 2017), leaf expansion rates and total leaf area (Monneveux *et al*., 2006; T. Liu *et al*., 2018). In maize, inadequate N supply impairs ultrastructure and function of chloroplast by reducing the number of chloroplasts per cell, lamellae per grana, which weakens photosynthesis and decreases biomass and grain yield (Jin *et al*., 2015; Z. Liu *et al*., 2018). The changes in leaf morphology and anatomy, as well as changes in chloroplast ultrastructure and the abundance of different photosynthetic pigments, that result from nitrogen deficit stress all alter the environment in which critical photosynthetic processes must function, including non-photochemical quenching (NPQ). The amount of light absorbed in plants often exceeds the capacity of the electron-transfer reactions (Pearcy, 1990; Burgess *et al*., 2016). Absorption of excess energy can result in over-reduction of photosystem II (PSII) antennas, leading to the production of damaging reactive oxygen species which can cause irreversible declines in photosynthetic efficiency (Müller *et al*., 2001). To prevent PSII from photodamage, plants employ NPQ to dissipate the excess energy absorbed by plant leaves. Faster NPQ induction and relaxation has been found to be associated with greater photosynthetic capacity and biomass in tobacco, soybean, and rice (Kromdijk *et al*., 2016; De Souza *et al*., 2022; Xin *et al*., 2023).

Previous studies have found extensive variation in the maximum level of NPQ observed in diverse populations of rice and Arabidopsis (Jung and Niyogi, 2009; Wang *et al*., 2017; Rungrat *et al*., 2019). We recently demonstrated that, under well fertilized field conditions over multiple years there is also significant, genetically controlled variation in the kinetics of how NPQ responses to sudden changes in light intensity which allowed us to identify distinct sets of genes controlling different attributes of the NPQ kinetics (Sahay *et al*., 2023*a*). However, comparatively little is known about how environmental perturbations, including nutrient deficits, impact natural variation in NPQ kinetics. We quantified NPQ kinetics at multiple developmental stages across a maize association panel grown in the field under low- and control-N. We identified diverse patterns of response/non-response to nitrogen treatment which were validated, for a subset of genotypes, across multiple additional environments and linked to differences in both CO_2_ assimilation and growth. We identify specific changes to the photosynthetic apparatus that are associated with, and may explain, why some maize genetics exhibit large changes in NPQ kinetics in response to nitrogen deficit stress while others do not.

## Materials and Methods

### Field experiments with the maize association panel

A collection of 225 diverse maize accessions (hereafter referred to as genotypes) from the Buckler-Goodman maize association panel (Flint-Garcia *et al*., 2005) were evaluated in a field experiment in 2019 (Table S1). A subset of 47 genotypes from the same population were also planted in the field in 2020. Both field experiments were conducted under non nitrogen limiting (hereafter referred to as control) and nitrogen limiting (hereafter referred to as low N) conditions at the University of Nebraska-Lincoln Havelock Research Farm, Lincoln, NE, USA. The 2019 field was located at N 40.853, W 96.611 and the 2020 field was located at 2020: N 40.859, W 96.598.

The 2019 field experiment has been previously described by (Meier *et al*., 2022) and (Rodene *et al*., 2022). Briefly, a completely randomized block (CRB) design experiment was set up with four blocks on 1^st^ June. Each block consisted of 252 plots with a complete set of 225 genotypes and a replicated genotype (B73×Mo17) included as a repeated check. Two blocks each were grown under control (54.43 kg/acre) and low N (no nitrogen applied) for a total of 1,008 plots. Replicates of the control and low N blocks were arranged diagonally, with the two control blocks in the North-East and South-West corners of the field and two low N blocks in North-West and South-East corners of the field. Each block was surrounded by border plots to minimize edge effects. Plots from the entire experiment were positioned in a grid of 176 (North-East and South-West) by 15 (North-West and South-East) plots (Table S2).

In 2020, the subset of 47 genotypes was planted on 22^th^ May in a CRB design in 8 blocks of 52 plots each. Each block included the complete set of 47 genotypes with B97 as a repeated check. Four blocks were assigned to control treatment and four blocks were assigned to low N treatment. A one plot (two rows) wide border was planted around the complete experiment while control and low N blocks were separated by a two-plot-wide border. In this study, NPQ data were collected from a subset of 23 genotypes among the 47 genotypes planted. The field was laid out in a grid of 18 (East-West) by 30 (North-South) plots. (Table S3).

In both years, plots consisted of two rows of the same genotype with a spacing of ∼75 centimeters between rows and ∼15 centimeters between sequential plants in the same row. In 2019, plots were 5.6-meter long, with 38 kernels planted per row. In 2020, plots were 1.5-meter long with 11 kernels planted per row. In both years, nitrogen fertilizer was applied in control plots in the form of urea prior to planting. No nitrogen was applied in low N plots. Standard agronomic practices were employed throughout the experiment to control the weeds.

### Plant propagation in growth chamber

The seeds were surface sterilized with 20% bleach for 20 min, followed by five times washing and overnight soaking in distilled water. Seeds were sown in 6-liter pots (13008000, Hummert International, Earth City, MO, USA) filled with Metromix 200 soil mixture (BM2 Germination and Propagation Mix; Berger, Saint-Modeste, Canada) in growth chambers (GR48, Conviron, Manitoba, Canada). Conditions in growth chambers were maintained at a 16-h photoperiod of 900 µmol m^-2^ s^-1^ light intensity, 25°C/20°C day/night temperature, and 65-70% relative humidity. In the preliminary experiment, four nitrogen treatments: control (210.2 mg of N L^-1^), moderate (140 mg of N L^-1^), low (105.1 mg of N L^-1^) and insufficient N (25 mg of N L^-1^) were tested on B73 maize genotype. After 4 days after germination, plants were irrigated with Hoagland media containing different nitrogen treatments twice a week, and soil was maintained with 90% field water capacity every day (for details about media composition see Supplementary Method 1). Based on the preliminary test on B73 maize genotype with four nitrogen treatments, low N treatment (105.1 mg of N L^-1^) was used for further experiments in the growth chamber due to a significant but moderate effect on physiological traits (*P* ≤ 0.04; Fig. 1A-G) and biomass (*P* ≤ 0.02; Fig. S1H). Oppositely, a small reduction in N (moderate N treatment i.e., 140 mg of N L-^1^) led only to no significant differences from control, however lowering N amount to 25 mg of N L^-1^ Hoagland media (insufficient N treatment) showed very early on yellowing leaf phenotype suggesting severe N deficiency. For the growth chamber condition experiments four genotypes were chosen from the maize association panel to represent increased (Group A genotypes; MS153 and Mo1W) or no-change (Group B genotypes; NC312, and H100) in *NPQ_max_* under low N conditions.

**Fig 1.**
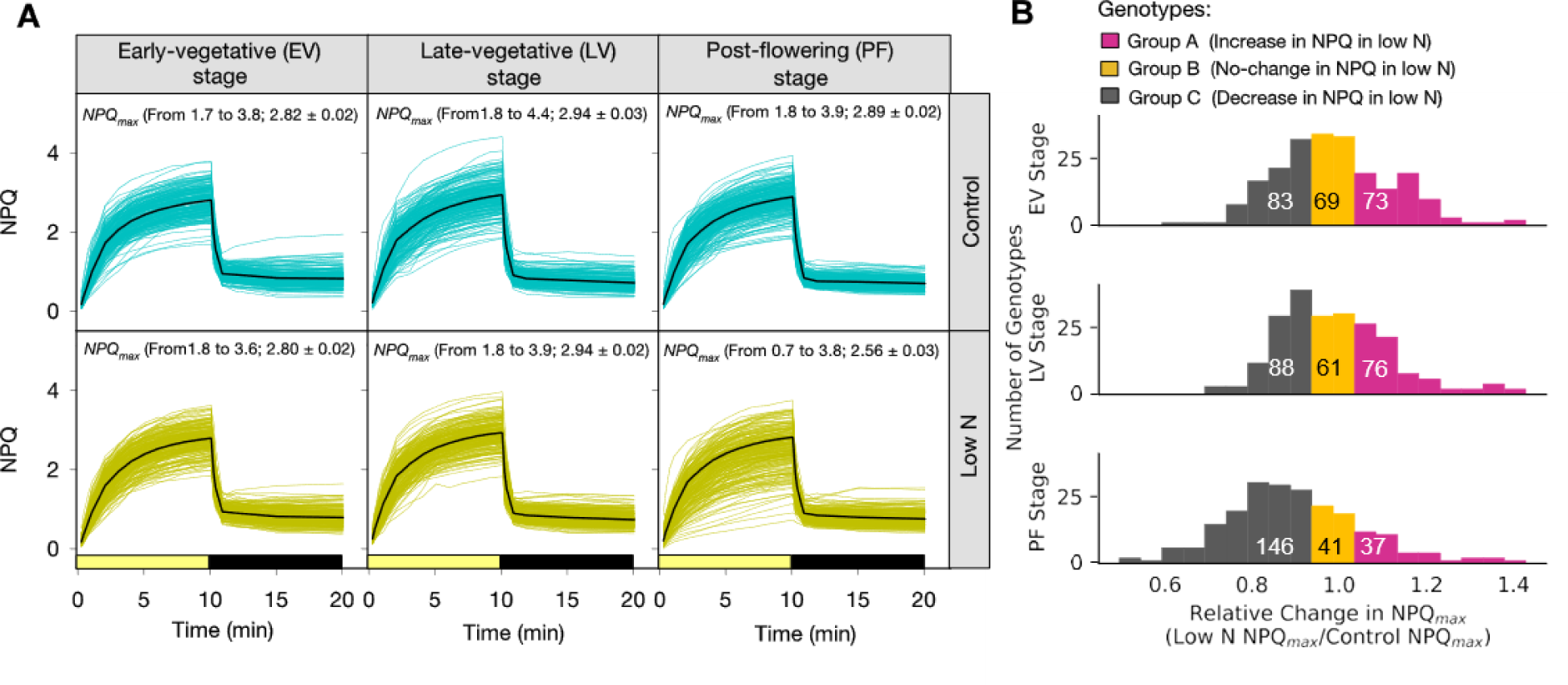
Distribution of kinetics curves of nonphotochemical quenching (NPQ) and maximum value of NPQ (*NPQ_max_*) at three developmental stages for maize (*Z. mays*) association panel grown in the field under control and low nitrogen conditions. **(A)** NPQ induction and relaxation kinetics for 225 genotypes at early-vegetative (EV), late-vegetative (LV), and 224 genotypes at post-flowering (PF) stages measured in 2019 under control and low N conditions. Each colored line represents the average value observed for an individual maize genotype across replicated plots of the same genotype in the same treatment and multiple leaf disks collected from each plot. Solid black lines indicate the overall mean NPQ response across all genotypes at a given developmental stage and treatment. The yellow horizontal bar indicates the timing of light treatment and the black horizontal bar represents the timing of the dark treatment. **(B)** Distribution of the relative changes in NPQ capacity of different maize genotypes under low nitrogen after 10 minutes of light treatment (*NPQ_max_*) in maize genotypes at each of the three developmental stages evaluated. The numbers over the gray, orange, and pink indicate number of genotypes exhibited a decrease (Group C; defined as ≤ 95% of value from the control conditions), no change (Group B), and increase (Group A; defined as ≥ 105% of corresponding control conditions) of the *NPQ_max_* under low N relatively to control conditions, respectively. Data used to produce this figure are given in Dataset 1.

### Measuring NPQ kinetics for field-grown and growth-chamber-grown maize plants

The NPQ kinetics was analyzed for the maize association panel grown in the field, 26 d, 47 d and 66 d after sowing in 2019, and 33 d, 49 d, and 76 d after sowing in 2020, that corresponded to early-vegetative (EV), late-vegetative (LV), and post-flowering (PF) stages, respectively. For each developmental stage, collection of data took four constitute days. In 2019 and 2020, two and three plants from the middle of the row were analyzed per plot, respectively. For the growth-chamber-grown plants the NPQ kinetics were measured 40 d after sowing plants corresponding to late vegetative stage. As previously described (Sahay et al., 2023), NPQ kinetics were investigated in semi-high-throughput manner on leaf disks (0.32 cm^2^) collected to the 96-well plate (781611; BrandTech Scientific, Essex, Ct, USA) from the sun exposed middle portion of the youngest fully expanded leaves. Leaf disks were cut using hand-held puncher between 16:00 and 18:30 h, positioned adaxial surface facing down into plate, covered with moist sponges and incubated overnight in the dark. Following day disks were imaged using a modulated chlorophyll fluorescence imager (FluorCam FC 800-C, Photon Systems Instruments, Drasov, Czech Republic). First the minimum (*F*_o_) and maximal (*F_m_*) fluorescence in the dark were imaged. Subsequently, the leaf disks was subjected to 10 min of 2000 μmol m^-2^ s^-1^ light (a combination of 1000 μmol m^-2^ s^-1^ of a red-orange light with λ_max_= 617 nm and 1000 μmol m^-2^ s^-1^ of a cool white 6500 K light) followed by 10 min of darkness. To capture changes in steady state fluorescence (*F*_s_) and maximum fluorescence under illuminated conditions (*F*_m_’) over time, the saturating flashes were provided at the following intervals (in s):15, 30, 30, 60, 60, 60, 60, 60, 60, 60, 60, 60, 60, 9, 15, 30, 60,180, and 300. Saturating flashes had intensity of 3200 μmol m^-2^ s^-1^ (a cool white 6500 K light) and a duration of 800 ms. For the growth-chamber-grown plants, in addition to the NPQ induction curve measured in 2000 μmol m^-2^ s^-1^ the NPQ was measured at four additional intensities of 150, 500, 900 and 4000 μmol m^-2^ s^-1^. NPQ was calculated using Eqn1, assuming the Stern– Volmer quenching model (Bilger and Björkman, 1994):

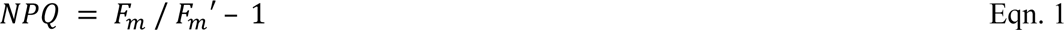

The NPQ measure here mostly represented two of the fastest components of NPQ named energy-dependent (qE) and zeaxanthin-dependent quenching (qZ).

Maximum PSII efficiency (*F_v_*/*F_m_*) was estimated using following equation (Genty *et al*., 1989):

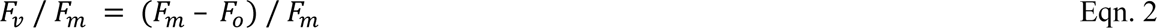

### Derivation of NPQ kinetic traits

The parameters attributed to rate, amplitude, and steady state of NPQ kinetics were obtained by fitting hyperbola and exponential equation to NPQ curves in the light (induction; Eqn. 3 and 4), or in the darkness (relaxation; Eqn. 5 and 6; Table S4) using MATLAB (Matlab R2019b; MathWorks, Natick, MA, USA). The complete list of the eight analyzed parameters and their biological meanings are presented in Table S4. Goodness of fit was defined as the discrepancy between measured values and the values predicted by the equation fit to the data.

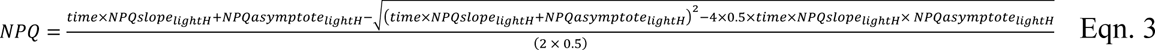

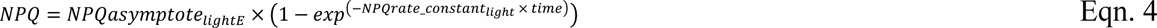

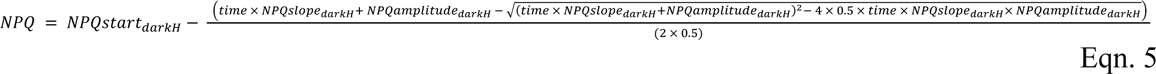

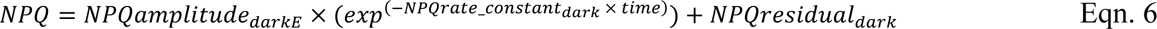

### Broad sense heritability and percent genetic variation

To analyze the effects of genotype, nitrogen condition, developmental stage, and their interactions on each trait, a linear model followed by analysis of variance (ANOVA) was used. The following model was fit:

Trait ∼ Genotype + Nitrogen_Treatment + Development_Stage + Genotype*\**Nitrogen_Treatment *+* Genotype*Development_Stage + Development_Stage*\**Nitrogen_Treatment *+ Genotype*\*Development_Stage*Nitrogen_Treatment

The sum of squares (SS) for each term in the model was extracted and divided by the total SS to obtain the percentage of variance explained by each term. To calculate broad-sense heritability, a separate linear model was fitted for each trait for each treatment condition and developmental stage. The variance due to genotype divided by total variance was used as a broad-sense heritability estimate.

### Fluorescence- and absorbance-based photosynthetic parameters

In 2020 field trial and growth-chamber experiments, fluorescence and absorbance-based parameters were measured on plants using spectrophotometers (MultispeQ V2.0; PhotosynQ, East Lansing, MI, USA; (Kuhlgert *et al*., 2016). All the measurements were performed using one of manufacture provided protocols, Photosynthesis Rides, that is based on Pulse-Amplitude-Modulation (PAM) method and MultispeQ data were analyzed with the PhotosynQ web application (https://photosynq.org). At late-vegetative stage of field (45-day-old) and growth chamber (35-days-old) grown plants, the transient value of theoretical NPQ (NPQ_T_) was estimated based on (Tietz *et al*., 2017):

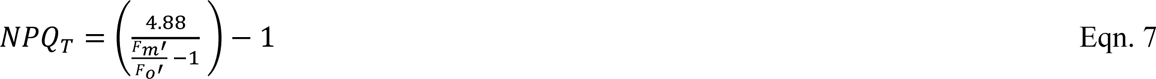

Fractions of light utilized to drive photochemical quenching (ϕPSII; named also PSII operating efficiency), dissipate as heat (ϕNPQ), and quenched by unregulated excitation dissipation (ϕNO) were calculated using Eqns. 8-10, respectively (Kuhlgert *et al*., 2016).

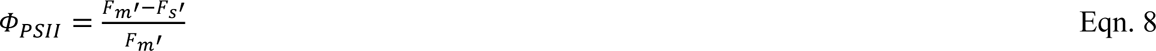

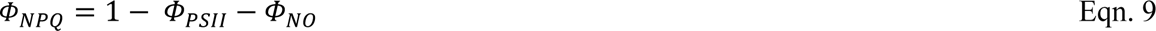

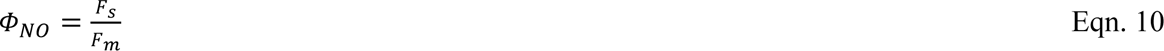

In addition, in growth-chamber experiment leaf thickness based on Hall Effect and leaf relative chlorophyll content (ChlPQ_SPAD_) based on absorbance at 650 and 940 nm were measured using MultispeQ (Mass et al., 1989; Kuhlgert *et al*., 2016). With the use of MutispeQ also electrochromic shift (ECS) based on Dark Interval Relaxation Kinetic (DRIK) method was used to monitor proton fluxes and the transthylakoid proton motive force (*pmf*) *in vivo*. The maximum amplitude of the ECS DIRK signal, related to the light-dark difference in thylakoid *pmf* was calculated according the Eqn 11 where tau (τ) is the relaxation time for the ECS signal, which is inversely related to the activity of the ATP synthase (Sacksteder *et al*., 2000).

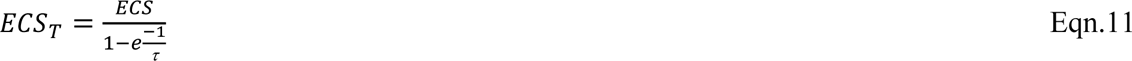

The proton conductivity (*g*H*^+^*) that is proportional to the aggregate conductivity of the thylakoid membrane to protons and largely dependent on the activity of ATP synthase was calculated after (Kanazawa and Kramer, 2002) using the Eqn. 12.

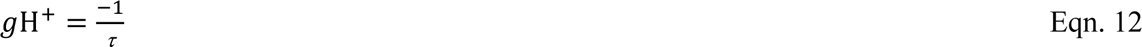

The redox state of photosystem I (PSI) reaction centers was estimated based on P700^+^ absorbance changes at 810 and 940 nm to monitor active centers (*PSI*_ac_), over-reduced center (*PSI*_orc_), and oxidized center (*PSI*_oxc_) after Klughammer and Schreiber (1994) and Kanazawa *et al*. (2017).

All the measurements were performed on the youngest fully expanded leaf. In the field, three randomly selected plants were measured per genotype from the middle of the row of each plot.

### Photosynthetic gas exchange and fluorescence measurements

Photosynthetic gas exchange measurements were performed on the youngest fully developed leaf of the growth-chamber-grown plants at late-vegetative stage (37 days after germination) using a LI-6800 equipped with an integrated modulated fluorometer (LI-COR, Inc. Lincoln, NE, USA). Pulse amplitude-modulated chlorophyll fluorescence measurements were used with the multiphase flash routine (Loriaux *et al*., 2013). The fluorescence was measured in parallel with gas exchange measurements at the range of light intensities to determine the light (Q) response of leaf net CO_2_ assimilation (*A*_n_), linear electron transport (*J*), photosystem II operating efficiency (ΦPSII) and NPQ. The [CO_2_] inside the cuvette, block temperature and water vapor pressure deficit were controlled at 400 µmol mol^-1^, 23°C, and 1.3 kPa, respectively. Leaf was dark adapted for 20 min in 6 cm^2^ cuvette after which *F*_o_ and *F*_m_ values were recorded to calculate *F_v_/F_m_* (Eqn. 2). For steady state measurements, light intensity was gradually increased from 0 to 25, 50, 75, 100, 125, 150, 200, 400, 600, 800, 1000, 1400, 1800 and 2000 µmol m^-2^ s^-1^. *A*_n_, stomatal conductance (*g*_s_), *F*_s_ and *F*_m_’ data were simultaneously recorded when reached steady state at each step which lasted between 300-600 s. For fluctuating light response measurements, the leaf first adapted to 2000 µmol m^-2^ s^-1^ to reach steady-state of gas exchange, then the light intensity was varied from 2000 to 1800, 1400, 1000, 800, 600, 400, 200, 150, 125, 100, 75, 50, and 25 µmol m^-2^ s^-1^. Each step lasted 4 min and was followed by 2000 µmol m^-2^ s^-1^ for 4 min. At each light intensity, gas exchange and fluorescence data were recorded trice every 80 s. The average values of three measurements were used to reconstruct the final light response curve.

For both light response curves *J* were calculated according to Eqn. 13.

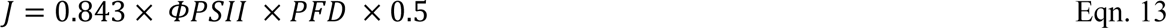

Quantum efficiency of CO_2_ assimilation (ΦCO_2max_) and quantum efficiency of linear electron transport (*ΦPSII*_max_) were derived from initial slope of the non-rectangular hyperbola equation fit to the light-response (*Q*) curves of *A*_n_ and *J*, respectively. *A*_sat_ and *J*_max_ were obtained from the asymptote of the non-rectangular hyperbola equation fit to *A*_n_/*Q* and *J*/*Q* curves, respectively.

To measure CO_2_ response to *A*_n_ (*A*/*C*_i_ curve), leaves were clumped in the cuvette in which light, [CO_2_], block temperature and water vapor pressure deficit were controlled at 2000 µmol m^-2^ s^-1^, 400 µmol mol^-1^, 23°C, and 1.3 kPa, respectively. After the leaf reached steady-state of gas exchange, the CO_2_ concentrations in the cuvette was varied from 400 to 50, 100, 150, 250, 350, 400, 500, 700, 900, and 1200 µmol mol^-1^. Gas exchange parameters were recorded when steady state was reached in each step. Non-rectangular hyperbola was fit to *A*_n_ response to intracellular CO_2_ concentration (*C*_i_) to obtain *in vivo* capacity for phosphoenol-pyruvate (PEP) carboxylation (*V*_pmax_; initial slope) and capacity for PEP regeneration and leakage of CO_2_ from the bundle sheath which determine the [CO_2_]-saturated rate of *A*_n_ (*V*_max_; asymptote of the curve) (von Caemmerer, 2000).

### Photosynthesis-related pigments quantification

After 42 d since sowing 0.635 cm^2^ or leaf tissue were collected from the youngest fully expanded leaf of the growth-chamber- and field-grown plants 4 h after the start of photoperiod. Tissue was grounded in liquid nitrogen and combined with 500 μl of pre-chilled 90% methanol (A452-1, Fisher Scientific, Hampton, NH, USA) vortexed and finally incubated for 4h on ice. The mixture was centrifuged for 5 min at 1902 × g at 4°C (5424R, Eppendrof, Enfield, CT, USA) and pellet washed with additional methanol until it became completely white. Supernatant was transferred to 96-well plates to measure the absorbance at 470, 652 and 665 nm using a plate reader (Microplate Reader HT, Bio Tek, Winooski, VT, USA). Chlorophyll (Chl) *a*, *b* and total carotenoid contents were determined using the measured absorptions and the equations of (Lichtenthaler, 1987).

### Analysis of growth and morphological traits

After 42 d since sowing the growth chamber growing plants were harvested to estimate leaf area, total fresh weight, and stalk height. The pictures of separate leaves were analyzed in ImageJ (National Institute of Health, Maryland, USA) to obtain a total leaf area. To evaluate the dry weight of above and below ground fraction of plants the roots, leaves and stalk were dried to constant weight at 60°C. In addition, the specific leaf area was calculated as the ratio of leaf area to leaf dry mass (Garnier *et al*., 2001).

### Statistical analysis

All statistical analysis of NPQ kinetics measurements was performed using SAS (version 9.4, SAS Institute Inc., Cary, NC, USA). Normal distribution and homogeneity of variance in data were tested with the Shapiro–Wilk and the Brown–Forsythe test, respectively. When either test rejected the null hypothesis, data was transformed and the tests were rerun. If one or both tests still rejected the null hypothesis the Wilcoxon non-parametric test was used in place of statistical tests which assume normal distributions. For data where both statistical tests failed to reject the null hypothesis, significant effects of treatment and genotype were evaluated using two-way ANOVA (α = 0.05) followed by Dunnett’s test or two-tailed Student t-test to address the comparison of low N treatment to control treatments. Accession means were used when Pearson correlation analysis was conducted for traits where measurements of the same genotype were made across multiple environments, mean values for each genotype in each treatment or stage were employed.

## Results

### Variation in NPQ kinetics across three developmental stages and two nitrogen treatments in 225 maize genotypes

A semi-high-throughput method based on fluorescence assay of leaf disks collected to 96-well plates (Sahay *et al*., 2023*a*) was used to quantify the NPQ kinetics in leaf disks collected from a field experiment consisting of a maize association panel grown under two nitrogen treatments with replication. In 2019, a combined total of 6,002 leaf disks were collected across three time points from a maize field experiment with replicated genotypes grown under both control and low nitrogen conditions. Data for 5,796 disks passed quality control and were employed in downstream analyses, providing information on the response of 225 diverse maize genotypes at the early vegetative (EV) and the late vegetative (LV) stage and 224 diverse maize genotypes at the post-flowering (PF) stages. Substantial variation in the kinetics of NPQ were observed among the genotypes at each developmental stage and under each treatment (Fig. 1A). The maximum NPQ over the 10-minute high-light period (*NPQ_max_*), one of the most commonly analyzed NPQ traits, showed a similar range from 1.7 to 4.4 (average of 2.9) at five of the six combinations of developmental stages and treatments (Fig. 1A). The exception was the PF stage under low N in which plants showed much lower values of *NPQ_max_* that extended substantially the range of this trait from 0.7 to 3.8 (average of 2.6). Based on change in *NPQ_max_* between treatments, maize genotypes were sorted into three groups: genotypes where *NPQ_max_* increased under low nitrogen conditions relative to control (Group A), genotypes where *NPQ_max_* did not change in response to nitrogen treatment (Group B) and genotypes where *NPQ_max_* decreased under low nitrogen conditions relative to control (Group C). At the early- and late-vegetative stages a similar fraction of genotypes belonged to each of these three groups. However, at the late-developmental stage when our nitrogen deficit treatment should produce the most severe stress, 64% of genotypes exhibited decreases in *NPQ_max_* under low-N-treatment (Fig. 1B).

In addition to *NPQ_max_* we also considered a set of six traits extracted from NPQ response curves that quantify the rate, steady state, and range of the NPQ response to light and dark in individual leaf samples. Significant differences between control and low N conditions for these NPQ kinetic traits were observed across the association panel at some but not all developmental stages (Fig. 2A). The average rate of NPQ induction and relaxation for the association panel (*NPQrate constant_light_* and *NPQrate constant_dark_*) were significantly different (*P* ≤ 0.01, two tailed t-test) between control and low N treatments at two and three developmental stages, respectively. The speed of NPQ relaxation in the dark was significantly faster under low N treatment than control conditions (*NPQrate constant_dark_*) at all three developmental stages. The effect of low-N-treatment on the speed of NPQ induction in the light (*NPQrate constant_light_*) was statistically significant, but not consistent across stages with a significant decrease and significant increase in early- and late-vegetative stages, respectively. Changes in NPQ steady state (*NPQasymptote_light_*, *NPQ_max_*) were only statistically significant at the post-flowering stage, where both traits were significantly lower in low-N than under control conditions. The post-flowering stage also exhibited significantly higher degrees of failure to completely relax NPQ at the end of dark incubation (10 minutes) (*NPQresidual_dark_*) under low N conditions relative to control.

**Fig 2.**
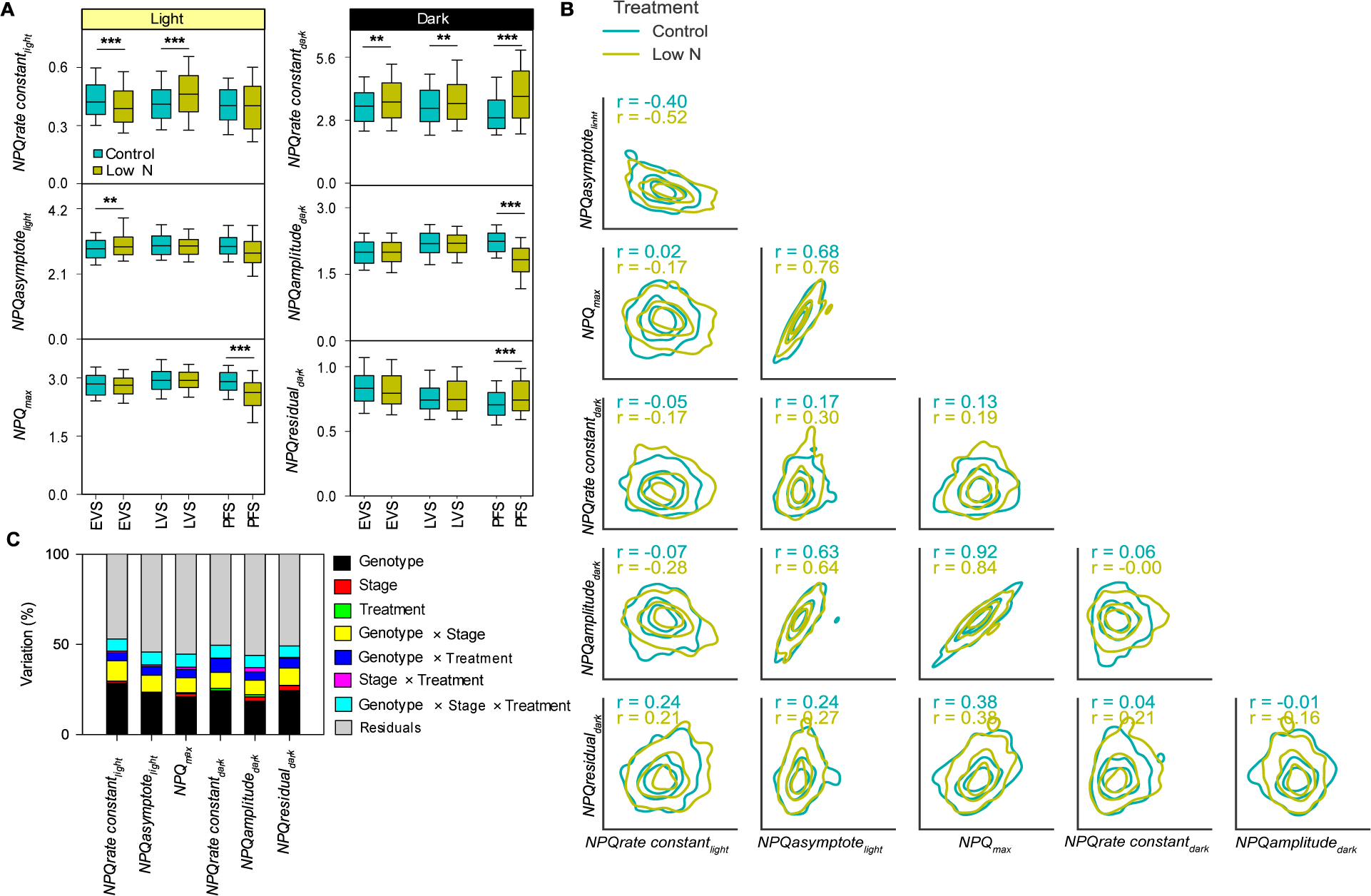
Variation in NPQ kinetic traits among N-treatments in three developmental stages and correlation between NPQ traits in late-vegetative stage for maize (*Z. mays*) association panel grown in the field. **(A)** Differences in the population level distributions of different traits describing specific attributes of NPQ kinetics. The speed of NPQ induction in the light and NPQ relaxation in the dark are described by *NPQrate constant_light_* and *NPQrate constant_dark_*, respectively. Steady state NPQ under high light and steady state NPQ in the dark after relaxation are described by *NPQasymptote_light_* and *NPQresidual_dark_*, respectively. The range of NPQ relaxation is described by *NPQamplitude_dark_*. *NPQ_max_* is the last value of NPQ in light (A full definition of each trait is provided in Table S4). The central line within each box plot indicates the median; the extent of the box indicates the 25th and 75th percentile values; upper and lower whiskers show either the minimum and maximum value or 1.5 × interquartile range, whichever results in a shorter whisker (n = from 224 to 225 genotypes). **(B)** Correlations among the six traits describing the attributes of NPQ kinetics at the late-vegetative stage. Contour lines in each plot indicate the density of maize genotypes for a pair of phenotypes under the two nitrogen treatments. Separate Pearson’s correlation coefficients (r) are displayed in each panel for each of the two N-treatments. **(C)** The percentage of variance explained by three factors (genotype, stage, treatment) and the percentage of variance explained by the interactions between these three factors for six traits describing attributes of NPQ kinetics for association panel grown in the field. In panel A and B asterisks indicate significant differences between control and low N (paired t-test; \**P ≤* 0.05; *\*\*P ≤* 0.01; *\*\*\*P ≤* 0.001). Data used to produce this figure are given in Dataset 1.

In all six combinations of N-treatment and developmental stage *NPQrate constant_light_* was significantly negatively correlated with *NPQasymptote_light_* and *NPQ_max_* (average Pearson correlation coefficient (r) = -0.46, *P* ≤ 0.001; Fig. 2B; S2 and S3). However, the rate of NPQ relaxation in the dark was only weakly correlated with the range of NPQ relaxation (average r = 0.06; *P* > 0.05) except at post-flowering stage in low N (r = 0.32; P ≤ 0.001). Several pairs of NPQ traits --*NPQrate constant_light_* & *NPQresidual_dark_* and *NPQ_max_* & *NPQresidual_dark_* --showed increased correlation at later developmental stages, independent of N treatment. Correlation between different NPQ kinetic traits tended to be higher in control conditions than in low N conditions. At the LV stage 11 out of 15 pairwise correlations were higher in low N conditions than in control (Fig. 2B). At the PF stage, a similar but weaker trend was observed with 10 out of 15 pairwise correlations between traits greater in low N than in control (Fig. S2B).

The proportion of variance explained by genotype -- as opposed to developmental stage, treatment or interaction terms -- was similar for all six NPQ kinetic traits, ranging from 19 to 29% of total variance (Fig. 2C). Treatment and developmental stage explained only minimal amounts of variance in the traits analyzed. However, the effect of interaction terms between genotype (G), developmental stage (S) and treatment (T) (G × S, G × T, S × T, and G × S ×T) ranged from 0.5 to 9.3% of total variance and collectively explained a similar portion of total variance to that explained by genotype (∼24%). When data from individual growth stages were analyzed independently, no one stage consistently exhibited greater broad sense heritability for NPQ kinetic traits averaging H^2^ = 0.47 (values ranged from 0.30 to 0.64) (Fig. S4).

### Consistent responses of NPQ to nitrogen treatment across years are associated with ratios of photosynthetic pigments

A subset of 23 maize genotypes selected based to represent all three patterns of response in *NPQ_max_* to low nitrogen were evaluated in a second field trial in 2020. These included 12 genotypes belonging to Group A (increased *NPQ_max_* under low N), 9 genotypes belonging to Group B (no change in *NPQ_max_* under low N) and 2 genotypes belonging to Group C (decrease in *NPQ_max_* under low N). At the late-vegetative stage a total 800 leaf disks were collected from the 2020 field experiment. Of these, 750 leaf disks passed quality control (QC) and were used for analysis. One genotype failed QC and was excluded, and a total 22 genotypes remained in the analysis. Twenty of the 22 remaining genotypes were placed into the same category, based on change in *NPQ_max_* between control and low N conditions, in 2020 as they had been in 2019 (Fig S5). The two exceptions were the genotypes -- IDS91 and NC332 -- placed in Group C (decrease in *NPQ_max_* under low N) in 2019, but which exhibited the same range of values as Group B (no change) genotypes in 2020. The *NPQ_max_* of these 22 genotypes in both years were correlated in both in control (Pearson’s correlation (r) = 0.44; *P* = 0.04) and low N conditions (r = 0.45; *P* = 0.03) (Fig 3A). Given the lack of consistent differences between Group B and Group C genotypes across multiple years, Group B and Group C genotypes are grouped together below. In order to determine whether the Group A/Group B differences were consistent throughout the day or varied diurnally, theoretical NPQ (NPQ_T_) was collected from plants in the field throughout the day (Fig. 3B). Group A genotypes exhibited significantly higher NPQ_T_ over most of the day (Fig. 3C; Fig. S6, S7). At the time point with highest light intensity (14:00 h) the NPQ_T_ was ∼1.8-fold higher (*P* < 0.001; Fig. 3C; Fig. S8) in Group A of genotypes on low N than on control. For Group B genotypes diurnal response of NPQ_T_ looked almost identical in both N treatments except for the last point of diurnal at 20:00 h. Group B genotypes also exhibited consistent quantum yield distribution into ϕ_NPQ_ (*P* = 0.14), ϕ_PSII_ (*P* = 0.29) and ϕ_NO_ (*P* = 0.07) across N treatments (Fig. S8), while Group A genotypes exhibited a significant shift towards a greater proportion of overall quantum yield being quenched by NPQ (1.2-fold; ϕ_NPQ;_ *P* = 0.0005) and a smaller proportion of overall quantum yield being captured by photosystem II (ϕ_PSII_) and unregulated excitation dissipation (ϕ_NO_; *P* = 0.001) (Fig. S8).

**Fig 3.**
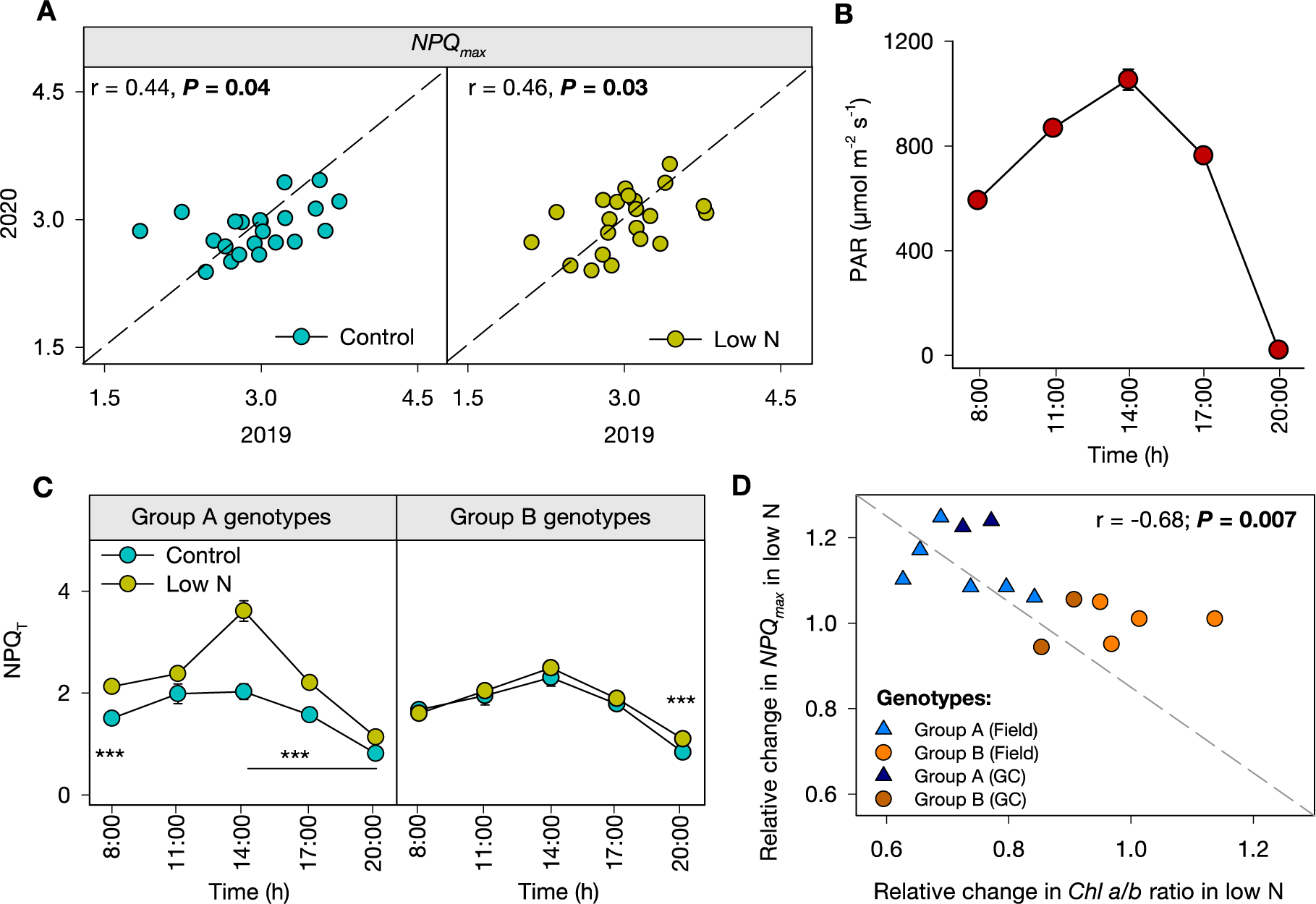
NPQ consistency across years, diurnal pattern and relation to chl *a*/chl *b* ratio in a subset of maize (*Zea mays* L.) genotypes in two nitrogen treatments at late-vegetative stage. **(A)** Correlation between the maximum levels of NPQ achieved after 10 minutes of illumination (*NPQ_max_*) for 22 maize genotypes in each of two field seasons under control and low nitrogen conditions. Each point corresponds to one of the 22 genotypes. X-axis and y-axis indicate the best linear unbiased predictor calculated for that trait for the same genotype in 2019 and 2020, respectively. Data are the means ± SEM (n = 4 plots, each plot mean is delivered from three biological replicates). **(B)** Diurnal photosynthetic active radiation (PAR) on 17^th^ July 2020. Data points are means ± SEM of control and low N together (n = 22 genotypes) at each time point but error bars frequently are smaller than the symbol size and not visible in these cases. **(C)** Theoretical NPQ (NPQ_T_) measured in Group A and Group B genotypes under control and low N conditions in field 2020 across a diurnal time course. Genotypes Group A – genotypes with upregulated NPQ in low N and Group B – genotypes that did not change in NPQ in low N in either 2019 or 2020. Data points are means ± SEM (n = 11 genotypes) but frequently are smaller than the symbol size. Asterisks indicate significant differences between control and low N based on two-tailed t-test (\*\*\**P ≤* 0.001). **(D)** Correlation between relative changes in Chl *a*/Chl *b* and *NPQ_max_* under low N in 10 genotypes grown in the field in 2020 and four genotypes grown in the growth chamber. Data points are means ± SEM (n = 4 plots in field; n = 4 plants in GC). In panel A and D, r and *P*-values correspond to Pearson’s correlation coefficient and the significance of tested correlations, respectively. *P*-values in bold are significant (*P ≤* 0.05). Data used to produce this figure are given in Dataset 2, 3 and 4.

Chlorophyll abundance was assayed in a subset of ten maize genotypes in the same field experiment. As expected, plants grown under low nitrogen treatment accumulated less total chlorophyll than did maize genotypes grown under control conditions. This reduction was proportionally greater for chlorophyll (Chl) *a* (average 18%; *P* < 0.0001) and proportionally smaller for Chl *b* (average 0.6%; *P* = 0.9) (Table 1). However, in Group B genotypes where *NPQ_max_* did not consistently change between nitrogen treatments, the ratio of Chl *a* to Chl *b* also remained largely constant between treatments. The greater decrease in Chl *a* under low N was driven entirely by Group A genotypes where the average Chl *a*/Chl *b* ratio declined from 3.1 under control conditions to 2.5 under low N conditions (average 18%; P ≤0.05). Consequently, the relative change in Chl *a*/Chl *b* and *NPQ_max_* on low N could be described by a single highly negative correlation (r = -0.68; P = 0.007; Fig. 3D). Group A genotypes also exhibited a significant drop in total carotenoid content (average ∼23%; *P* ≤ 0.04) which was not observed in Group B genotypes (*P* ≥ 0.33).

**Table 1.**
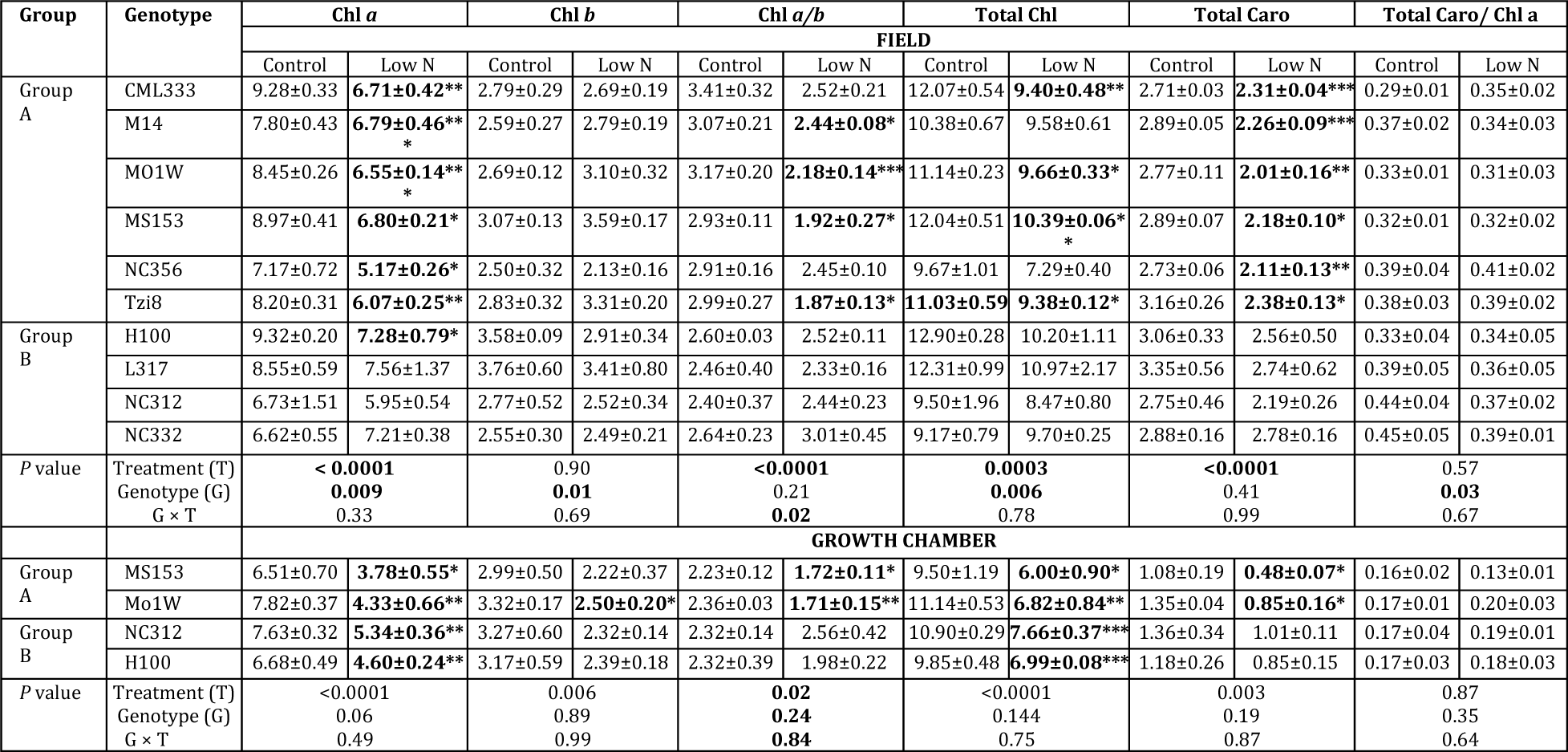
Quantification of photosynthesis-related pigments in Group A (increase NPQ in low N) and Group B (no-change in NPQ in low N) genotypes under control and low nitrogen treatments in the field and growth chamber.

### Maize genotypes with elevated NPQ under low N exhibit compromised photosynthesis and reduced biomass accumulation

The responses of two pairs of genotypes representing Group A, increased NPQ under low N (MS153 and Mo1W) and Group B, no change in NPQ under low N (NC312 and H100) were also evaluated in controlled environment (e.g. growth chamber) conditions. Differences in NPQ kinetics between nitrogen treatments were largely consistent between growth chamber and field trial data at the late vegetative stage (Fig. 4A) and these same four genotypes exhibited largely consistent NPQ kinetic responses to low nitrogen in multiple years and at multiple development stages (S9 and S10). Majority of NPQ traits were higher in Group A genotypes under low N relative to control at the late vegetative stage (Fig. 4B) while Group B genotypes showed no statistically significant change in NPQ traits between treatments (∼ 0.95%; *P* ≥ 0.32). The increase in NPQ induction under low nitrogen conditions among Group A genotypes remained statistically significant when NPQ was induced by 900 or 2000 µmol m^-2^ s^-1^ light (*P* ≤ 0.01) and Group B genotypes showed no change in NPQ induction in response to low N (*P* ≥ 0.08), however at higher (4000 µmol m^-2^ s^-1^) or lower (150 and 500 µmol m^-2^ s^-1^) light intensities there were not consistent differences between groups (Fig. S11A-E). Patterns of change in Chl *a*/Chl *b* ratios between treatments in the growth chamber were also consistent with the larger field study: Group A genotypes exhibited significant decreased in Chl *a*/Chl *b* ratios under low N (*P* ≤ 0.02) while Group B genotypes exhibited consistent (P ≥ 0.48) Chl a/Chl b ratios between low N and control conditions (Table 1).

**Fig 4.**
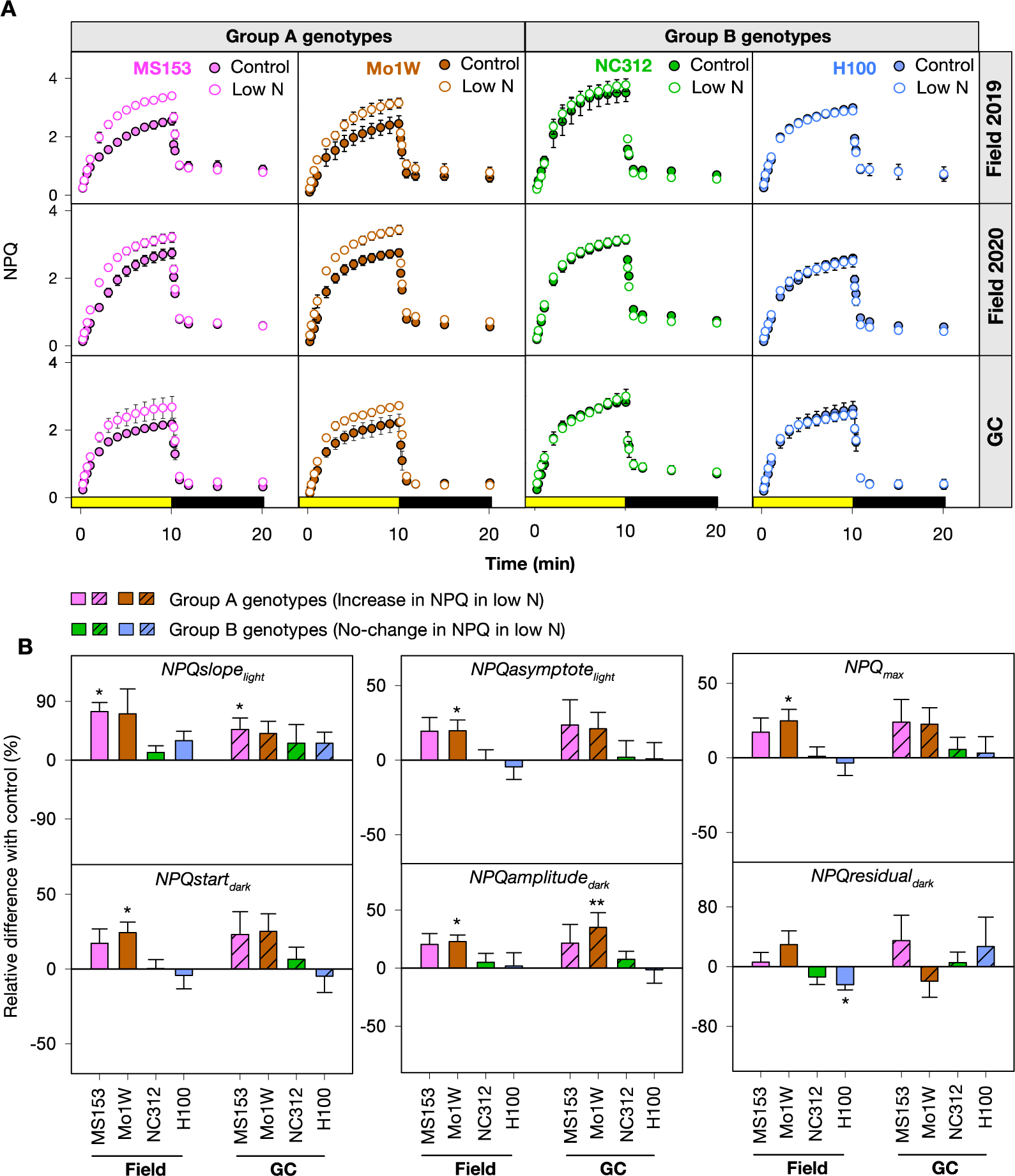
Comparison of NPQ kinetic responses to nitrogen treatments across three environments in four maize (*Zea mays* L.) genotypes at the late-vegetative stage. **(A)** Response of NPQ induction in light (indicated in yellow horizontal bar) followed by relaxation in dark (indicated by black horizontal bar) in two groups of genotypes; MS153, Mo1W (increase NPQ in low N; Group A genotypes), and NC312 and H100 (no change in NPQ in low N; Group B genotypes) in the 2019 and 2020 field seasons and in growth chamber (GC). **(B)** Relative differences (%) in each NPQ trait in light (*NPQslope_light_, NPQasymptote_light_,* and *NPQ_max_*) and dark (*NPQstart_dark_*, *NPQamplitude_dark_*, and *NPQresidual_dark_*) for each genotype under low N correspond to their control treatments in 2020 field and GC conditions. Symbols and bars are the means ± SEM (n = 4 plots in field and n = 4 plants in GC). In panel B, asterisks indicate significant differences between control and low N based on two paired t-test (\**P ≤* 0.05; *\*\*P ≤* 0.01; *\*\*\* P ≤* 0.001). Data used to produce this figure are given in Dataset 1, 2 and 5.

Group A genotypes showed an increase in proton motive force (ECS_T_) under low N, as estimated from nondestructive spectrometric and fluorescence measurements, similar in magnitude to the increase in NPQ_T,_ (∼56%; Fig. S12A and B). Group B genotypes also showed an increase in ECS_T_ under low N but the increase was smaller than that observed for Group A. All genotypes showed increases in the fraction of PSI oxidized centers (*PSI_oxc_*) and decreases both in proton conductivity (*g*H*^+^*) and fraction of photosystem I active centers (*PSI_ac_*) under low N relative to control, however these changes tended to be larger in magnitude in Group A than in Group B (Fig. S12 C, D and F).

Group A genotypes had on average higher net CO_2_ assimilation (*A*_n_), photosynthetic electron transport (*J*) and stomatal conductance (*g_s_*) than Group B genotypes under control conditions (Fig. 5). However, Group A genotypes lost relatively more photosynthetic capacity particularly under low N at high light intensities. For instance, *A*_sat_ decreased by 75% (*P* ≤ 0.0005) in both MS153 and Mo1W. However, in NC312 and H100, the same trait decreased but only by 25% (*P* = 0.002) and 10% (*P* = 0.28), respectively (Fig. 5F). Genotypes showed very similar differences in *J*_max_ to these observed for *A*_sat_ (Fig. 5H). In the light-limited portion of the *A_n_*/light-response (*Q*) and *J*/*Q* curves the difference between treatments and genotypes were less pronounced. Nevertheless, quantum efficiency of leaf net CO_2_ assimilation (*ΦCO_2max_*) decreased by an average of 29% (*P* ≤ 0.02) in Group A genotypes and only a statistically insignificant 14% (P ≤ 0.8) in Group B genotypes (Fig. 5G). PSII operating efficiency (*ΦPSII_max_*) followed a similar trend to that observed for *ΦCO_2max_* (Fig. 5I). The preceding results were measured under constant light, however results under fluctuating light were largely consistent with those under constant light (Fig. S13) with the exception of the *g_s_*/*Q* curves (Fig. S13C) where stomatal response to low light was reduced under low N relative to control. CO_2_ response curves (*A*/*C*_i_ curves) showed substantial variation among genotypes in the maximal rate of phosphoenolpyruvate (PEP) carboxylation (*V_pmax_*) and [CO_2_]-saturated rate of *A* (*V_max_*) determined by capacity for PEP regeneration and leakage of CO_2_ from the bundle sheath were not significantly different (*P* > 0.05) between treatments for any of the four analyzed genotypes (Fig. 5G-O).

**Fig 5.**
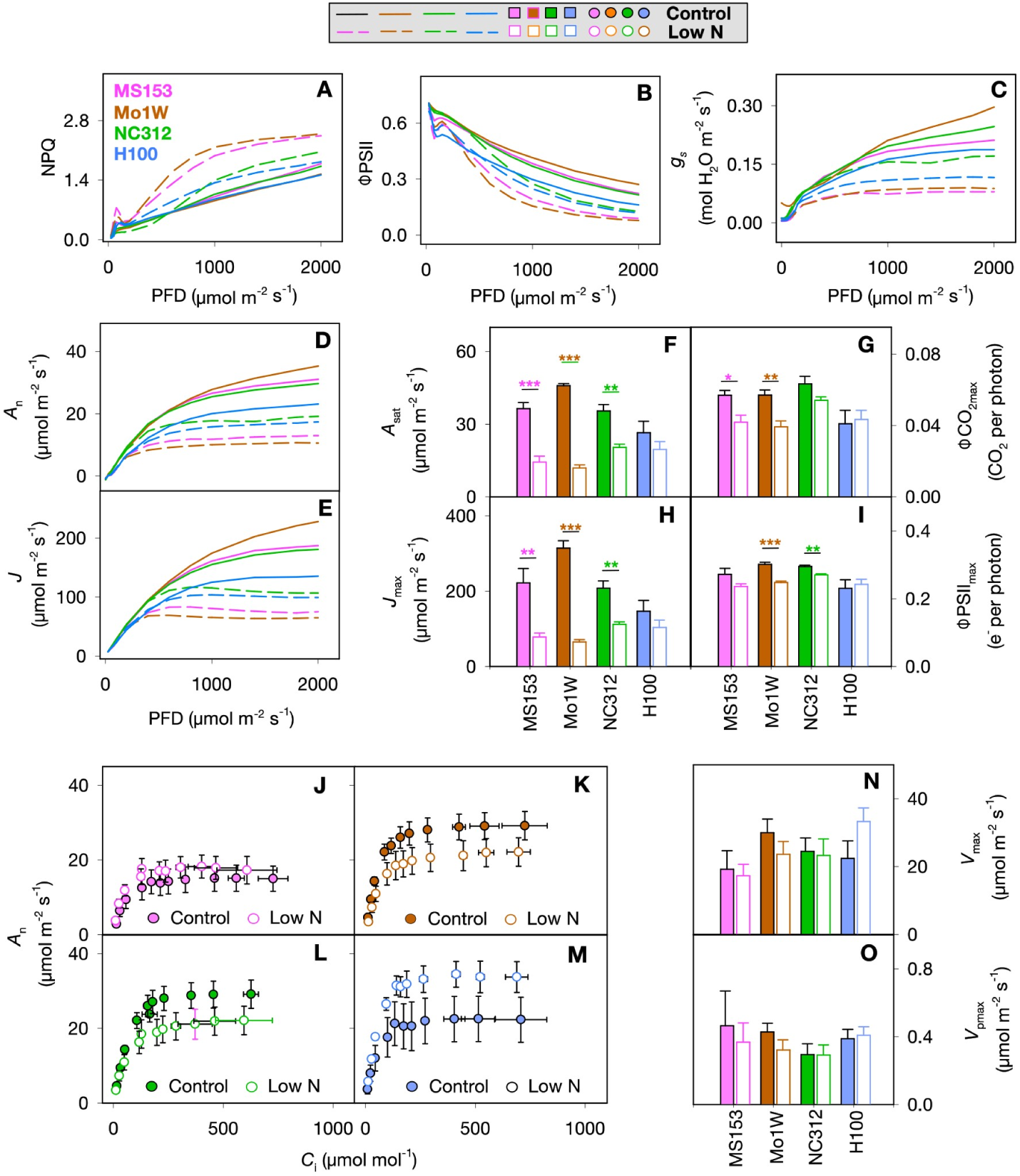
Response of gas exchange and fluorescence related photosynthesis parameters to steady-state changes in light and intercellular CO_2_ concentration (C_i_) in four maize (*Zea mays* L.) genotypes grown under two nitrogen treatments at growth-chamber conditions. **(A)** NPQ, **(B)** photosystem II operating efficiency (*ΦPSII*), **(C)** stomatal conductance (*g*_s_), **(D)** net CO2 fixation rate (*A*_n_), and **(E)** linear electron transport rate (*J*) as a function of step increase in incident photon flux density (PFD). **(F)** Quantum efficiency of leaf net CO_2_ assimilation (*ΦCO_2max_*), **(G)** light-saturated rate of net CO_2_ assimilation rate (*A*_sat_), **(H)** maximal linear electron transport rate (*J*_max_), and **(I)** quantum efficiency of linear electron transport (*ΦPSII_max_*). (J-M) *A*_n_ measured as a function of gradual changes in *C*_i_. **(N)** Maximal velocity of Rubisco for carboxylation (*V*_pmax_), and **(O)** [CO_2_]-saturated velocity of *A*_n_ (*V*_max_). All traits were measured in the late-vegetative stage in MS153, Mo1W (increase NPQ in low N; Group A genotypes), and NC312 and H100 (no-change in NPQ in low N; Group B genotypes) under control (solid line, bar and circle) and low N (dashed line, open bar or open circle) treatments. Data are the means ± SEM (n = 4 biological replicates). Asterisks show significant differences in low N from the control (paired t-test; \**P ≤* 0.05; *\*\*P ≤* 0.01; *\*\*\* P ≤* 0.001). Data used to produce this figure are given in Dataset 6 and 7.

The changes in NPQ kinetics, photosynthetic pigment ratios, and photosynthetic traits between Group A and Group B genotypes under low N were associated with different biomass accumulation outcomes. All four genotypes showed substantial decrease in plant size on low N relative to control conditions, however, the decreases in biomass were proportionally greater for Group A genotypes than Group B genotypes (Fig. 6). Low N treatment resulting in a reduction in total fresh weight, above ground dry biomass, and root dry mass of ∼35% each among the two Group A genotypes (*P* ≤ 0.004, *P* ≤ 0.04, and *P* = 0.0005) (Fig. 6A) as well as a reduction in leaf area of ∼25% (*P* ≤ 0.03). The same traits in Group B genotypes were reduced by ∼15% with only total fresh weight and leaf area being significantly lower in one out of two analyzed genotypes (NC312; *P* ≤ 0.007).

**Fig 6.**
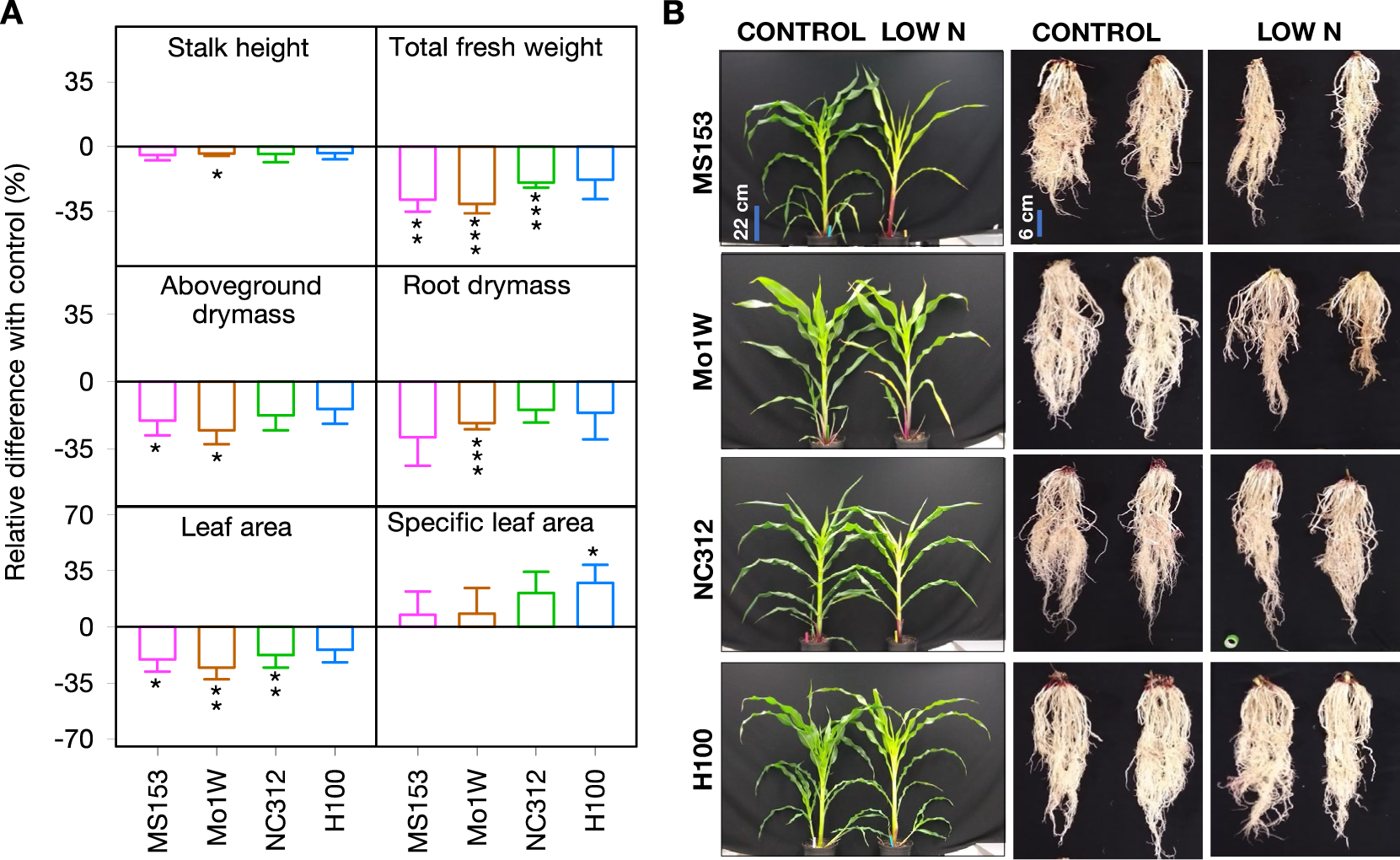
Morphological responses of four maize (*Zea mays* L.) genotypes to control and low nitrogen treatments. **(A)** Stalk height, total fresh weight, above-ground drymass, root dry mass, leaf area and specific leaf area under low N expressed as relative difference with the same traits measured from the same genotypes under control conditions (%). **(B**) Illustration of changes in above- and below- ground plant organs between treatments. All traits were measured in the late-vegetative stage for all four genotypes -- MS153, Mo1W (Group A), and NC312 and H100 (Group B) -- under control and low nitrogen treatments. Above-ground scale bar = 22 cm and below-ground scale bar = 6 cm. Data are the means ± SEM (n = 4 biological replicates). Asterisks indicate significant differences between low nitrogen and control (paired t-test; \**P ≤* 0.05; *\*\*P ≤* 0.01; *\*\*\* P ≤* 0.001). Data used to produce this figure are given in Dataset 8.

## Discussion

The importance of NPQ as a key determinant of photosynthetic productivity in natural systems, where plants are typically exposed to rapid changes in light intensity, has been appreciated for many decades (Demmig-Adams and Adams, 1992; Murchie and Niyogi, 2011; Kaiser *et al*., 2018; Wang *et al*., 2020). As a core component of photosynthesis and hence plant fitness, the activity of NPQ might be expected to be conserved within species. However a growing body of evidence supports the existence of substantial genetically controlled variation in the maximum potential to induce NPQ within species ranging from arabidopsis (Rungrat *et al*., 2019) to rice (Jung and Niyogi, 2009; Wang *et al*., 2017; Rungrat *et al*., 2019)) and soybean (Burgess *et al*., 2020). In the diverse panel of 225 maize genotypes we evaluated here, the difference in *NPQ_max_* between the genotypes most and least responsive to high light treatment was >2x in each combination of treatment and developmental stage (Fig. 1A). This degree of phenotypic diversity was approximately double the range reported for the same trait in surveys of diverse arabidopsis (Rungrat *et al*., 2019) and soybean (Burgess *et al*., 2020) populations. In the low nitrogen treatment at the post-flowering stage of development, the stage at which the effects of low nitrogen stress became most extreme, this range expanded even further to a 5x difference between the genotypes most and least responsive to high light treatment.

The extension of the study of NPQ kinetics from studying single genotypes to populations has enabled the identification of both substantial genetically controlled diversity, and specific genetic loci controlling variation in NPQ kinetic traits (Wang *et al*., 2017; Rungrat *et al*., 2019; Sahay *et al*., 2023*a*). However, optimal NPQ kinetic responses to maximize photosynthetic productivity likely vary across environments. In our initial study, we were able to identify sets of maize genotypes where *NPQ_max_* increased, decreased, or remained constant under low N treatment relative to control conditions (Fig. 1B). Of these three potential patterns, genotypes exhibiting two potential responses (increased *NPQ_max_* under low N and no change in *NPQ_max_* under low N) continued to exhibit consistent responses across multiple years as well as between field and controlled environment conditions (Fig. 3A; Fig. 4). Higher ECS_T_ (estimates proton motive force) and decreased *g*H*^+^*(estimates ATPase synthase activity) in thylakoid membrane in genotypes where NPQ increased in response to nitrogen deficit may reflect the activation of NPQ (Fig. S12) via protonation of the photosystem II subunit S protein (Li *et al*., 2000; Kanazawa and Kramer, 2002; Rott *et al*., 2011; Kanazawa *et al*., 2017; Takagi *et al*., 2017). Similar dependence of NPQ upregulation to downregulation of chloroplastic ATP synthesis was observed in cold-treated tobacco (Yang *et al*., 2018).

While total chlorophyll (except NC332) and carotenoid content declined in all maize genotypes tested under low nitrogen conditions, significant shifts in Chl *a*/Chl *b* ratios was observed only in the set of maize genotypes in which *NPQ_max_* increased significantly under low nitrogen treatment relative to controls (Tabel 1). This result is consistent with a previous report that an increase in *NPQ_max_* was associated with a decrease in Chl *a*/Chl *b* ratios in a chinese maize hybrid when grown under low N (Lu and Zhang, 2000). Similar observations have also been reported for cassava (*Manihot esculenta*) grown on low N (Cruz *et al*., 2003). Interestingly, in other stress related to nutrients availability (salt stress) in sorghum (*Sorghum bicolor*; (Netondo *et al*., 2004) and beach plum (*Prunus maritima*; (Zai *et al*., 2012) preserved similar Chl *a*/Chl *b* to that in control what correlated with unchanged NPQ.

The significant relative decrease in chlorophyll *a* suggests a greater loss in reaction center complexes and reengineering of the photosynthetic apparatus among maize genotypes where *NPQ_max_* increases under low N treatment. That means that relatively more excitation energy could be funneled to fewer PSII reaction centers in the responsive maize genotypes than the unresponsive ones. This increase in excitation energy per reaction center under low N may explain why an upregulation of NPQ is required to maintain adequate level of photoprotection in these genotypes, while not being required in maize genotypes which preserve similar ratios of Chl *a*/Chl *b* across both treatments.

Our results also suggest that the increase in NPQ activation under low N was insufficient to compensate for the deleterious effects of low N. Carbon assimilation (*A*_sat_) decreased more in genotypes where NPQ increased in response to low N conditions than in genotypes where NPQ remained unchanged which may explain the decreases in stomatal conductance (*g*_s_) which were also observed in these genotypes. Despite the difference in NPQ response to low N between genotypes, there were no differences in *F_v_/F_m_* between control and low N in pre-flowering and late-vegetative stages suggesting reduced carbon assimilation was not the result of photoinhibition (Fig. S14). While modest reductions in PEP carboxylation rate and Rubisco regeneration activity were observed in all four genotypes, these changes were not statistically significant and insufficient to explain the reduction in overall carbon assimilation. Instead, the decrease in carbon assimilation appears to be directly linked to the change in NPQ, in turn, Chl *a*/Chl *b* ratios which, among genotypes where NPQ increases in response to low nitrogen, results in a smaller proportion of light energy being captured by photosystem II (*ΦPSII*) and reduced electron transport (*J*_max_) relative to non-responsive genotypes. This result, like the change in Chl *a*/Chl *b* ratios would be consistent with a greater decrease in photosynthetic reaction centers in genotypes where NPQ responses to low nitrogen stress. As might be expected, these changes in photosynthetic activity were associated with substantially bigger declines in fresh and dry biomass for responsive maize genotypes under low N relative to control conditions (Fig. 6), although these results should be interpreted with caution given the relatively small numbers of responsive and non-responsive genotypes evaluated.

NPQ kinetics have long been known to change in response to the broadly defined environmental factors (Harvaux and Kloppstech, 2001; Fernández-Marín *et al*., 2021; Nosalewicz *et al*., 2022; Rodrigues de Queiroz *et al*., 2023). More recently the kinetics of NPQ have also been shown to vary as a result of naturally occurring genetic diversity in a range of crop and wild species (Wang *et al*., 2017; Rungrat *et al*., 2019; Burgess *et al*., 2020; Sahay *et al*., 2023*a*,*b*). Here we have demonstrated that genetic and environmentally controlled variations are not independent of each other. Instead, different genotypes exhibit differential plasticity of NPQ response to the same environmental perturbations. These genetically controlled differences in NPQ responses to environmental changes are, in turn, associated with differences in both the photosynthetic apparatus and photosynthetic productivity, as well as differences in overall biomass accumulation. The existence of naturally occurring genetic variation controlling NPQ trait plasticity increases the likelihood it will be possible to optimize NPQ kinetics in crop plants for different environments. Our results also suggest that large increases in NPQ may serve as a comparatively low cost and high throughput method to identify crop genotypes where photosynthetic health and productivity is compromised in different environments or in response to different stresses, particularly as we were able to recapitulate the differences between responsive and nonresponsive genotypes initially observed from leaf disk assays with in-field estimates of NPQ (e.g. NPQ_T_).

## Supporting data

Supporting figures, tables, methods and datasets described in this study are available in the online version of this article.

**Fig. S1.**
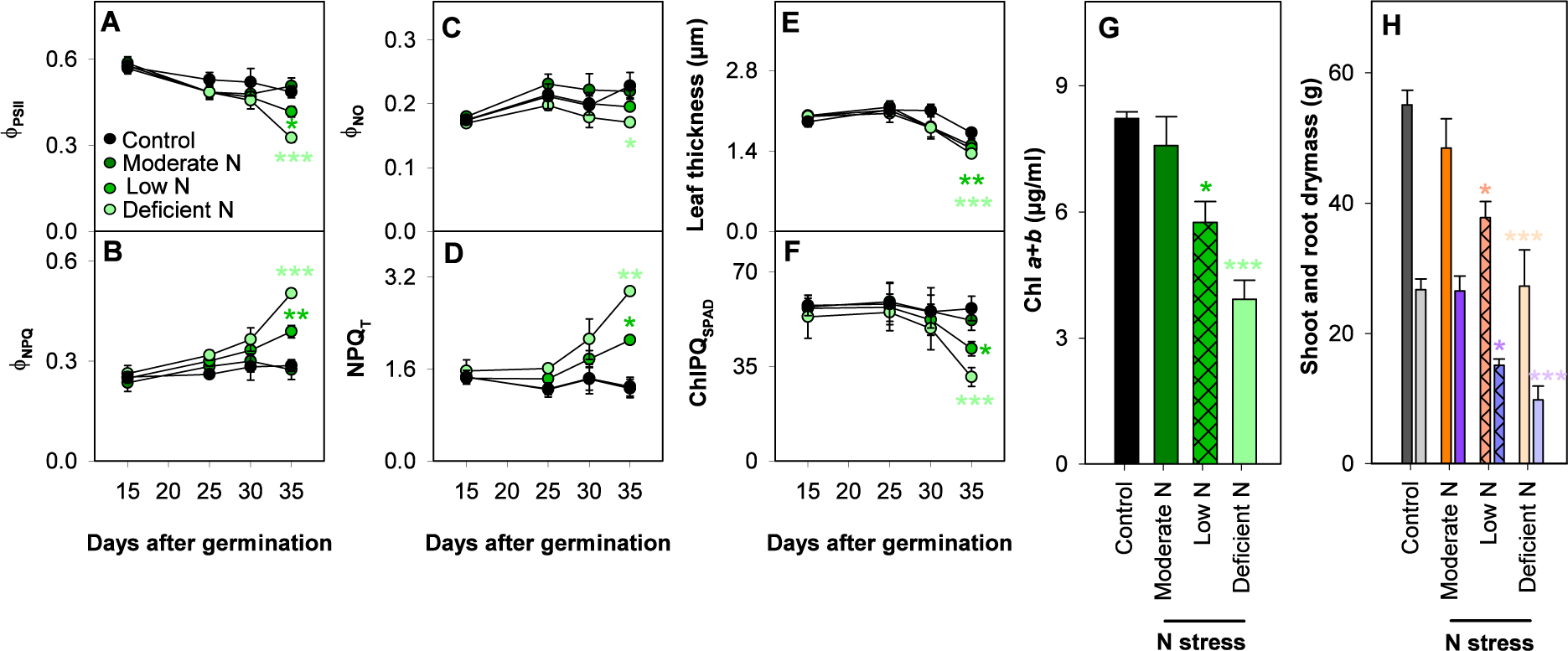
Evaluation of four nitrogen treatments in the maize reference genotype (B73) in the growth chamber. In panels A-F changes in physiological traits, and biomass are shown as a function of days after germinations in control, moderate, low, and deficient N conditions. The 15- to 35-day-old plants of maize genotype B73 were used to measure **(A)** quantum yield of photosystem II (Ф_PSII_), **(B)** quantum yield of NPQ (Ф_NPQ_), **(C)** quantum yield of unregulated non-photochemical losses (Ф_NO_), **(D)** theoretical NPQ (NPQ_T_), **(E)** leaf thickness, **(F)** relative total chlorophyll content estimated via SPAD method S (ChlPQ_SPAD_), **(G)** Leaf total chlorophyll *a* and *b* content measured in leaf extract. In addition, for 35-day-old plants **(H)** the total drymass of plants was measured. Data are the means ± SEM (n = 4 biological replicates). Asterisks show significant differences from the control (Dunnett’s two-way test; \**P* ≤ 0.05; \*\**P* ≤ 0.01; \*\*\**P* ≤ 0.001). Data used to produce this figure are given in Dataset 9.

**Fig S2.**
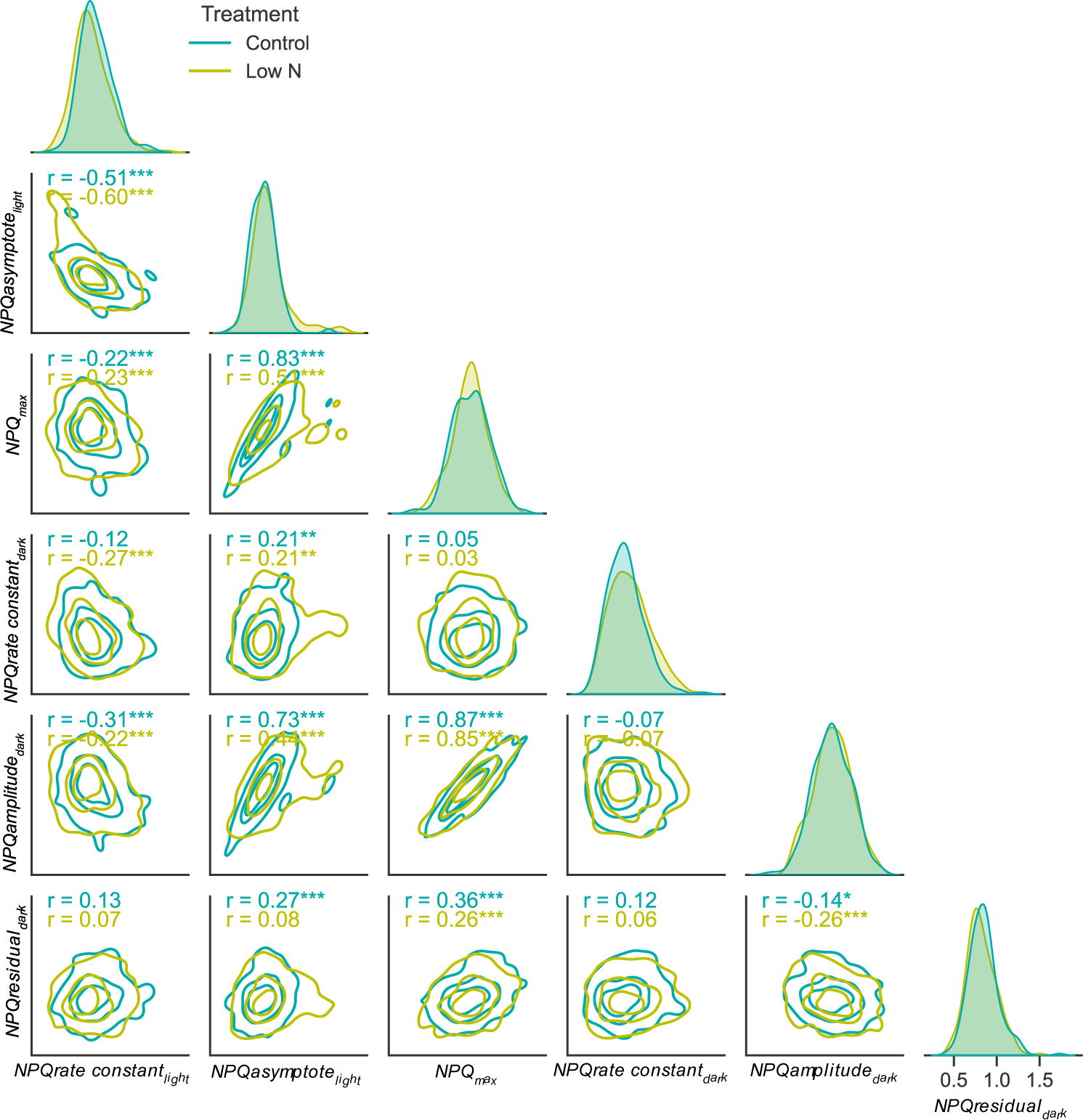
Correlation among six NPQ kinetic traits across the maize (*Zea mays* L.) diversity panel under control and low N treatments at early-vegetative stage. Each plot uses contour lines to mark the two-dimensional kernel density of maize genotypes for a pair of phenotypes under two nitrogen (N) treatments. Rate of NPQ induction and relaxation is described by *NPQrate constant_light_* and *NPQrate constant_dark_*, respectively. Steady state of NPQ induction and relaxation is described by *NPQasymptote_light_* and *NPQresidual_dark_*, respectively. The range of NPQ relaxation is described by *NPQamplitude_dark_*. The *NPQ_max_* is the last value of NPQ in light (a full definition of each trait is provided in Table S4). Pearson’s correlation coefficients (r) are displayed in each panel and asterisks indicate significant differences between control and low N inside the developmental stage (paired t-test; \**P* ≤ 0.05; \*\**P* ≤ 0.01; \*\*\**P* ≤ 0.001). Data corresponds to NPQ kinetics as shown in Figure 1A. Data used to produce this figure are given in Dataset 1.

**Fig S3.**
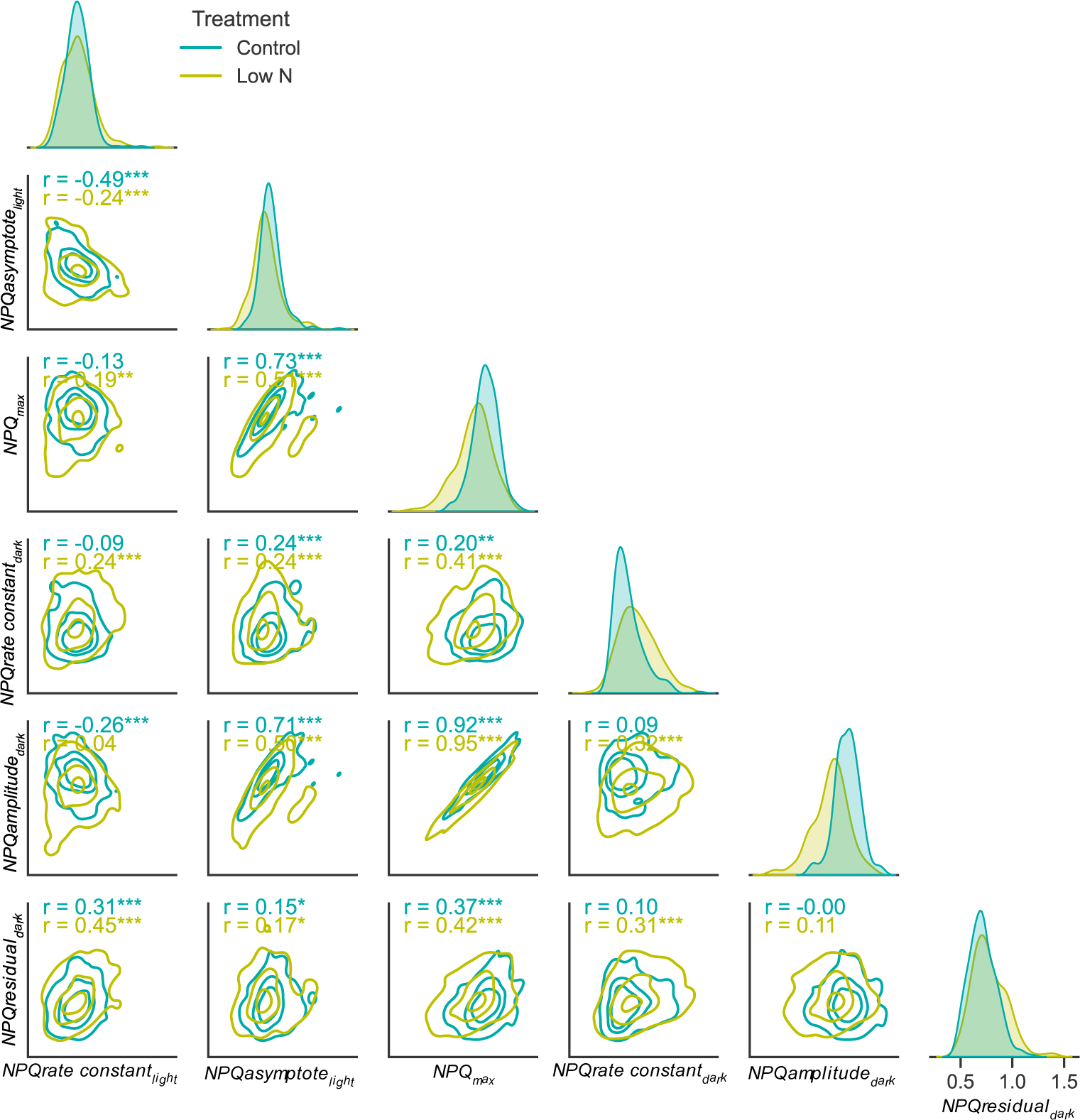
Correlation among six NPQ kinetic traits across the maize (*Zea mays* L.) diversity panel under control and low N treatments at post-flowering stage. Each plot uses contour lines to mark the two-dimensional kernel density of maize genotypes for a pair of phenotypes under two nitrogen (N) treatments. Rate of NPQ induction and relaxation is described by *NPQrate constant_light_* and *NPQrate constant_dark_*, respectively. Steady state of NPQ induction and relaxation is described by *NPQasymptote_light_* and *NPQresidual_dark_*, respectively. The range of NPQ relaxation is described by *NPQamplitude_dark_*. The *NPQ_max_* is the last value of NPQ in light (a full definition of each trait is provided in Table S4). Pearson’s correlation coefficients (r) are displayed in each panel and asterisks indicate significant differences between control and low N inside the developmental stage (paired t-test; \**P* ≤ 0.05; \*\**P* ≤ 0.01; \*\*\**P* ≤ 0.001). Data corresponds to NPQ kinetics as shown in Figure 1A. Data used to produce this figure are given in Dataset 1.

**Fig S4.**
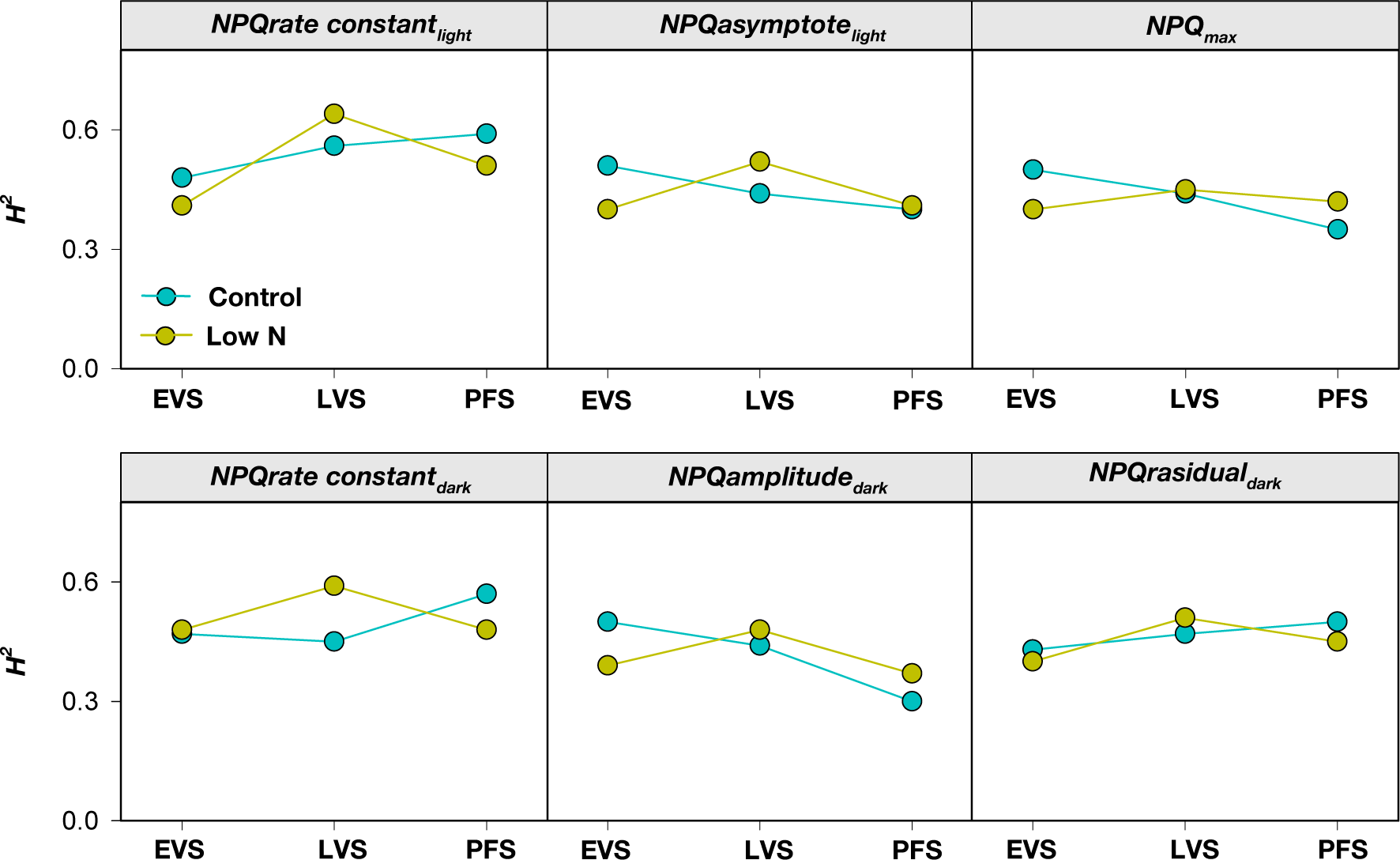
Broad sense heritability (*H_2_*) of six NPQ kinetic traits in maize (*Zea mays* L.) at three developmental stages and two nitrogen treatments. Rate of NPQ induction and relaxation is described by *NPQrate constant_light_* and *NPQrate constant_dark_*, respectively. Steady state of NPQ induction and relaxation is described by *NPQasymptote_light_* and *NPQresidual_dark_*, respectively. The range of NPQ relaxation is described by *NPQamplitude_dark_*. The *NPQ_max_* is the last value of NPQ in light (a full definition of each trait is provided in Table S4). Data corresponds to NPQ kinetics as shown in Figure 1A. Each data point showing heritability values estimated from 225 genotypes at early-vegetative (EV), late-vegetative (LV), and post-flowering (PF) stage under control and low N, except 224 genotypes at PF stage in low N condition in field 2019. Data used to produce this figure are given in Dataset 1.

**Fig S5.**
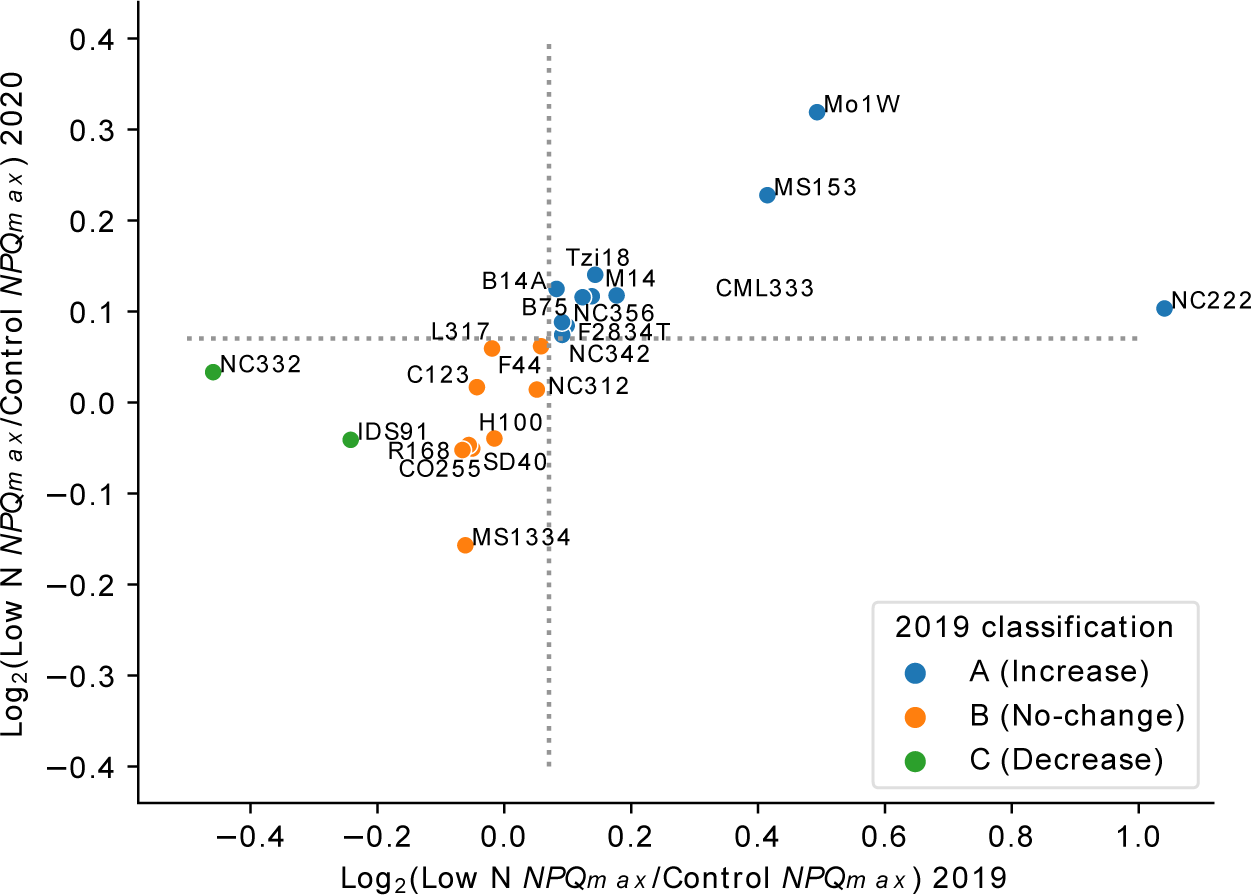
Distribution of 22 maize (*Zea mays* L.) genotypes according to classification to one of three possible groups of *NPQ_max_* evaluated in both 2019 and 2020. The three groups are: increase (Group A), no change (Group B), and decrease (Group C) of *NPQ_max_* in low N. Dotted lines indicate the threshold for a genotype to be classified as Group A vs Group B in 2019 (vertical line) and 2020 (horizontal line). Data used to produce this figure are given in Dataset 2.

**Fig S6.**
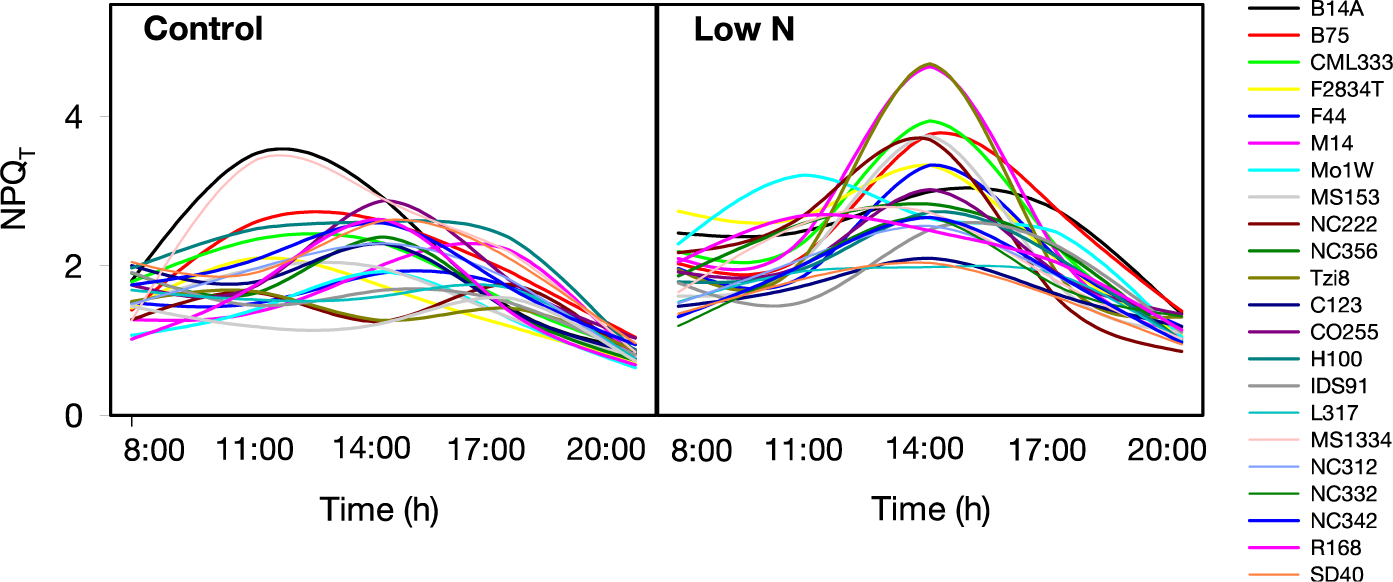
Diurnal variation in theoretical NPQ (NPQ_T_) in 22 maize (*Zea mays* L.) genotypes under control and low nitrogen in 2020 field trial. NPQ**_T_** was measured at 8:00, 11:00, 14:00, 17:00 and 20:00 h. The lines represent the mean trend of diurnal response for individual genotypes (n = 4 plots). Data used to produce this figure are given in Dataset 3.

**Fig. S7.**
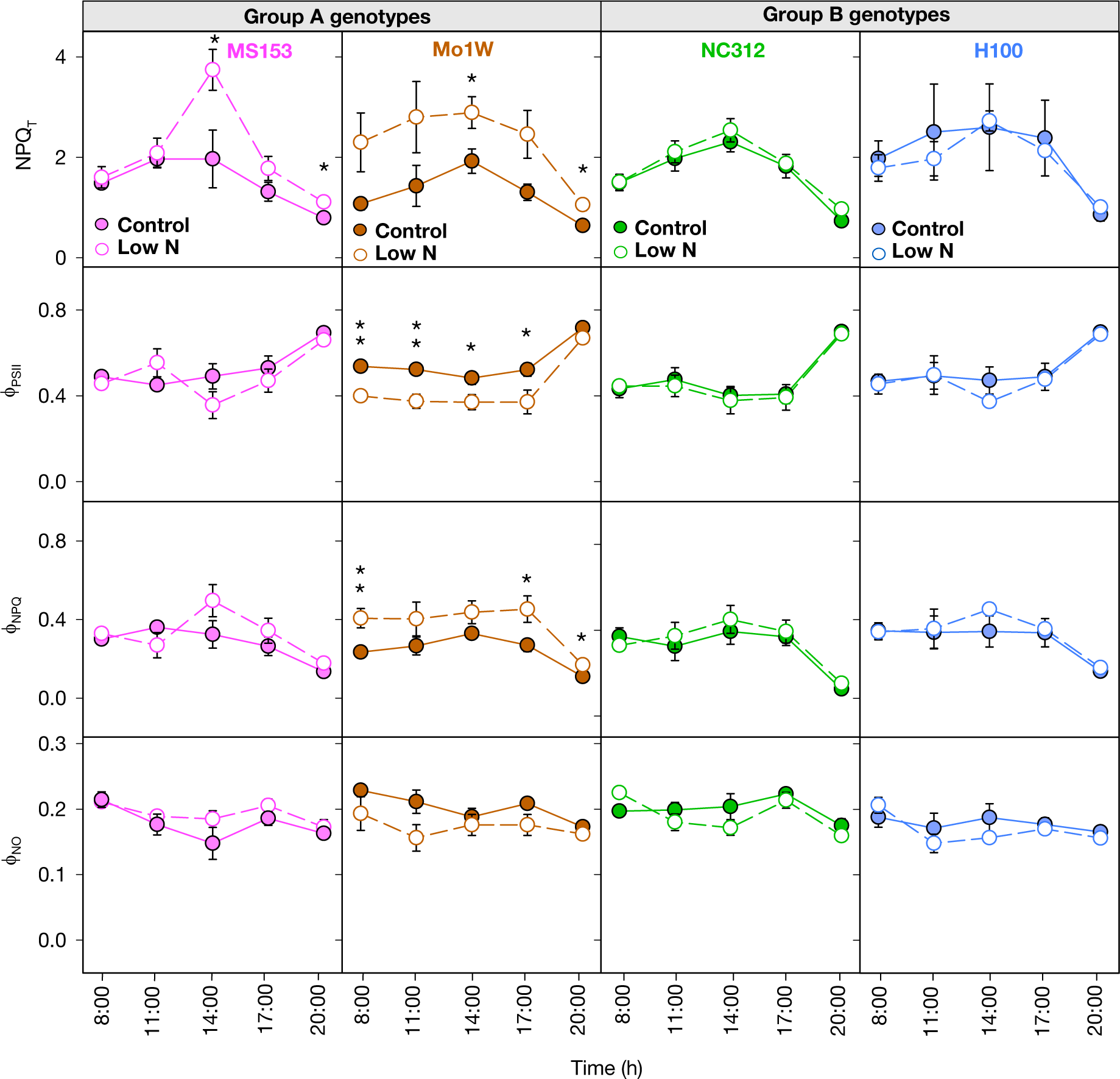
Diurnal profile of fluorescence-delivered traits in four selected maize (*Zea mays* L.) genotypes grown under two nitrogen treatments in 2020 field trial. Traits were measured in the late-vegetative stage in MS153, Mo1W (increase in NPQ in low N; group A genotypes), and NC312 and H100 (no-change in NPQ in low N; group B genotypes). Theoretical NPQ (NPQ_T_); quantum yield of photosystem II (Ф_PSII_); quantum yield of NPQ (Ф_NPQ_), and quantum yield of unregulated non-photochemical losses (Ф_NO_) are shown as a function of time of measurements. Symbols represent means ± SEM (n = 4 plots). Asterisks indicate significant differences between control and low N based on two-tailed t-test (\**P* ≤ 0.05; \*\**P* ≤ 0.01). Data used to produce this figure are given in Dataset 3.

**Fig. S8.**
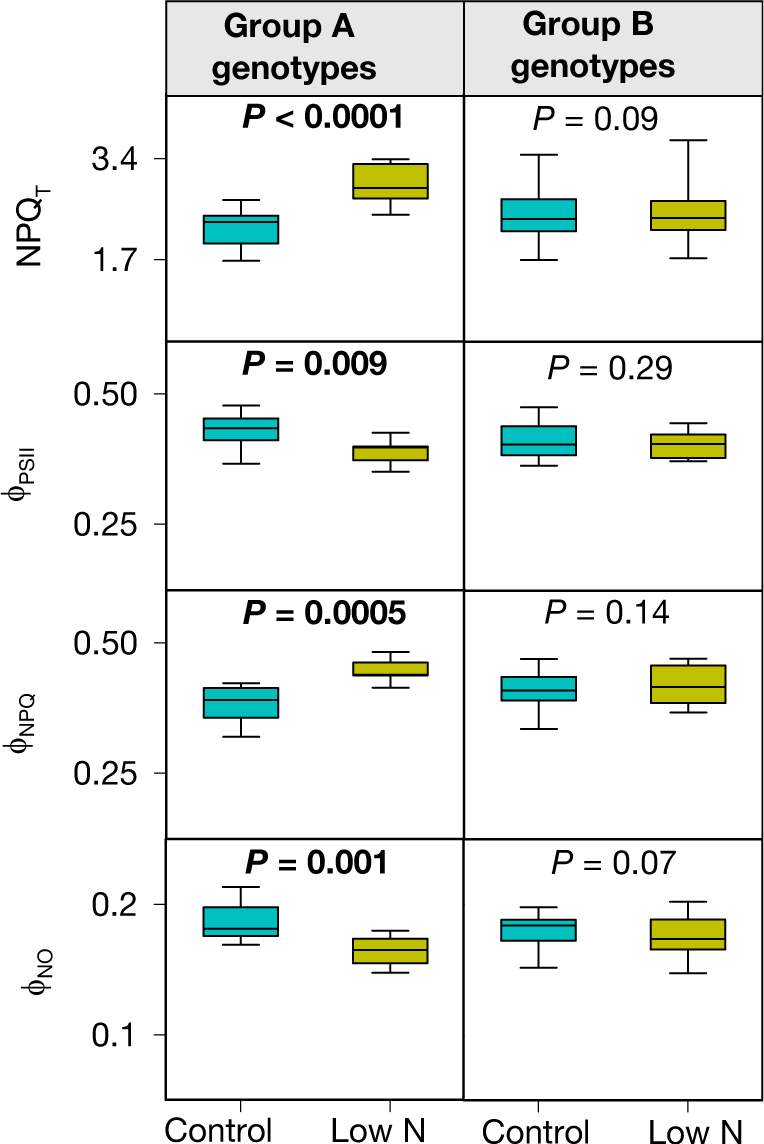
Response of fluorescence-delivered traits in group A (increase of NPQ in low N) and group B (no-change in NPQ in low N) maize (*Zea mays* L.) genotypes grown at low and control nitrogen in the field trial in 2020. **(A)** Theoretical NPQ (NPQ_T_). (**B)** Quantum yield of photosystem II (ϕ_PSII_). (**C)** Quantum yield of NPQ (ϕ_NPQ_). (**D)** Quantum yield of unregulated non-photochemical losses (ϕ_NO_). Eleven genotypes represented each group. Box plots represent a range of variations with the central line within each box plot representing the median; box edges indicate the upper and lower quantiles; upper and lower whiskers show minimum and maximum at the 1.5 × interquartile range. *P*-values were determined using a two-way ANOVA test and represent an effect of treatments (). Data used to produce this figure are given in Dataset 10.

**Fig S9.**
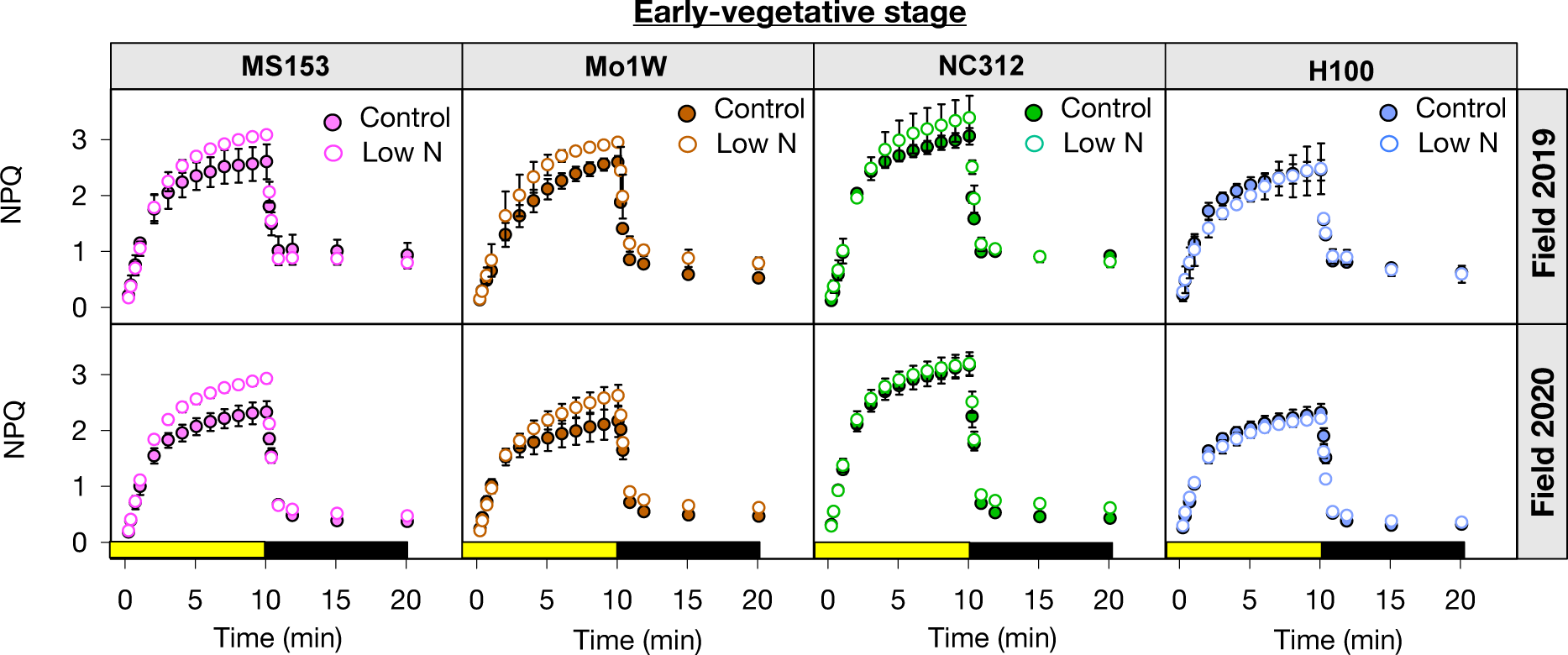
Comparison of NPQ kinetic responses to nitrogen treatments across two environments in maize (*Zea mays* L.) at the early vegetative stage. Response of NPQ induction in light (indicated in yellow horizontal bar) followed by relaxation in dark (indicated by black horizontal bar) in two genotypes MS153, Mo1W (increase NPQ in low N; group A genotypes), and NC312 and H100 (no change in NPQ in low N; Group B genotypes) in the 2019 and 2020 field trials. Symbols are the means ± SEM (n = 4 plots). Data used to produce this figure are given in Dataset 1 and 2.

**Fig S10.**
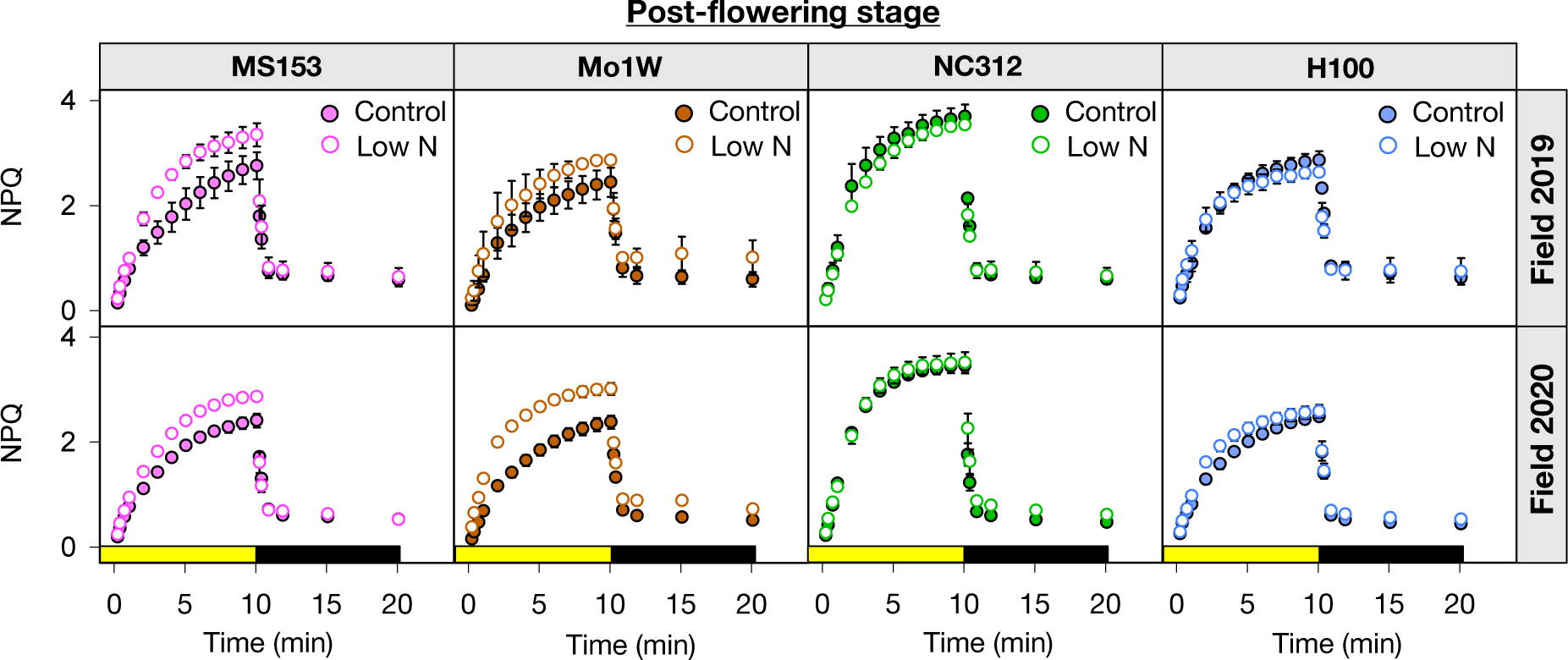
Comparison of NPQ kinetic responses to nitrogen treatments across two environments in maize (*Zea mays* L.) at the post-flowering stage. Response of NPQ induction in light (indicated in yellow horizontal bar) followed by relaxation in dark (indicated by black horizontal bar) in two genotypes MS153, Mo1W (increase NPQ in low N; group A genotypes), and NC312 and H100 (no change in NPQ in low N; Group B genotypes) in the 2019 and 2020 field trials. Symbols and bars are the means ± SEM (n = 4 plots). Data used to produce this figure are given in Dataset 1 and 2.

**Fig S11.**
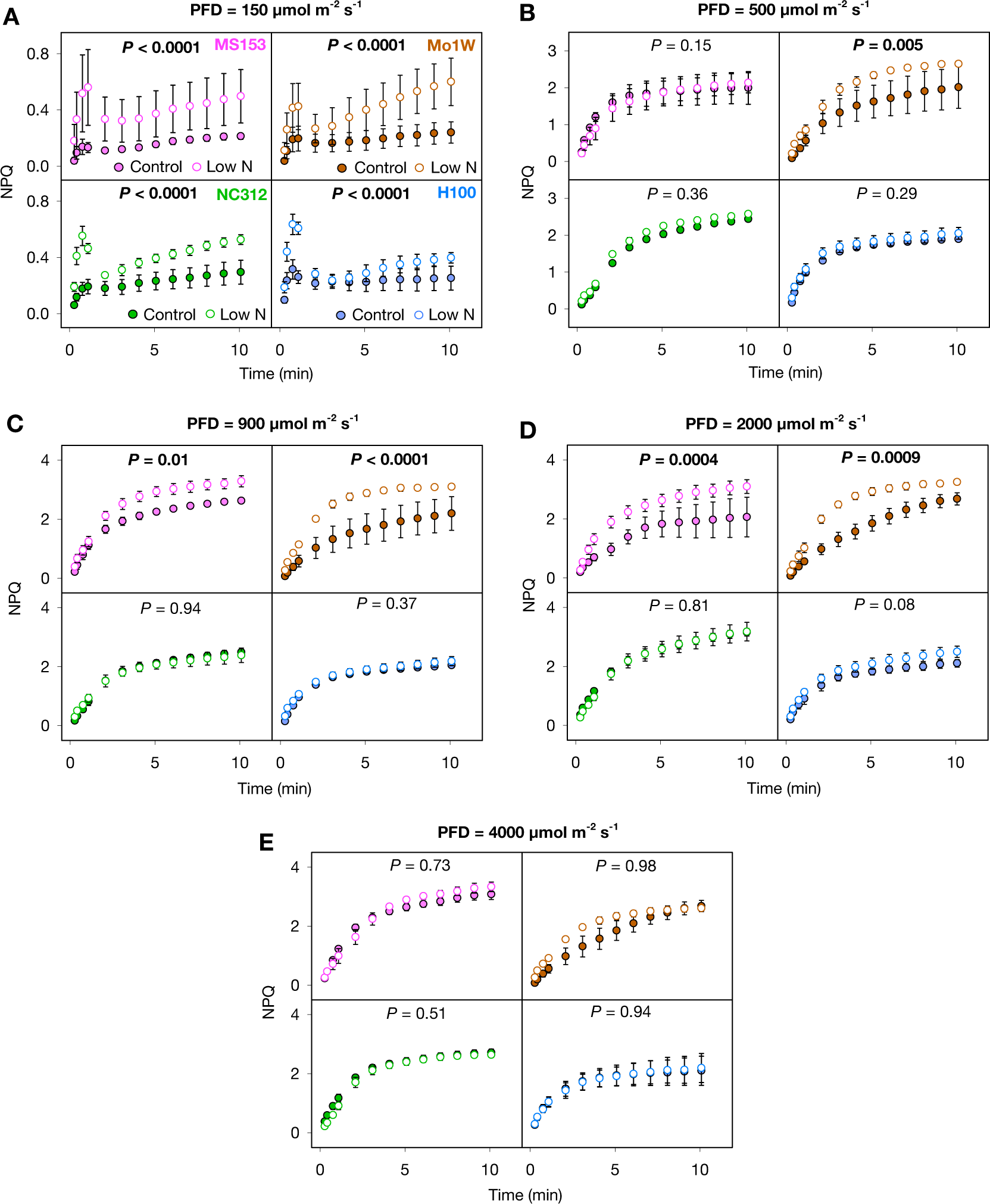
NPQ kinetics at five light intensities in four selected maize (*Zea mays* L.) genotypes grown under two N treatments in the growth chamber. Response of NPQ induction during 10 min illumination of (**A)** 150, (**B),** 500 (**C)** 900, (**D)** 2000 and (**E)** and 4000 µmol m^-2^ s^-1^ of photons flux density (PFD) in MS153, Mo1W (increase NPQ in low N; Group A genotypes), and in NC312 and H100 (no change in NPQ in low N; Group B genotypes) under control and low N. Data represents means ± SEM (n = 4 plants). *P*-values determine the effect of N-treatment in a two-way ANOVA. Data used to produce this figure are given in Dataset 11.

**Fig S12.**
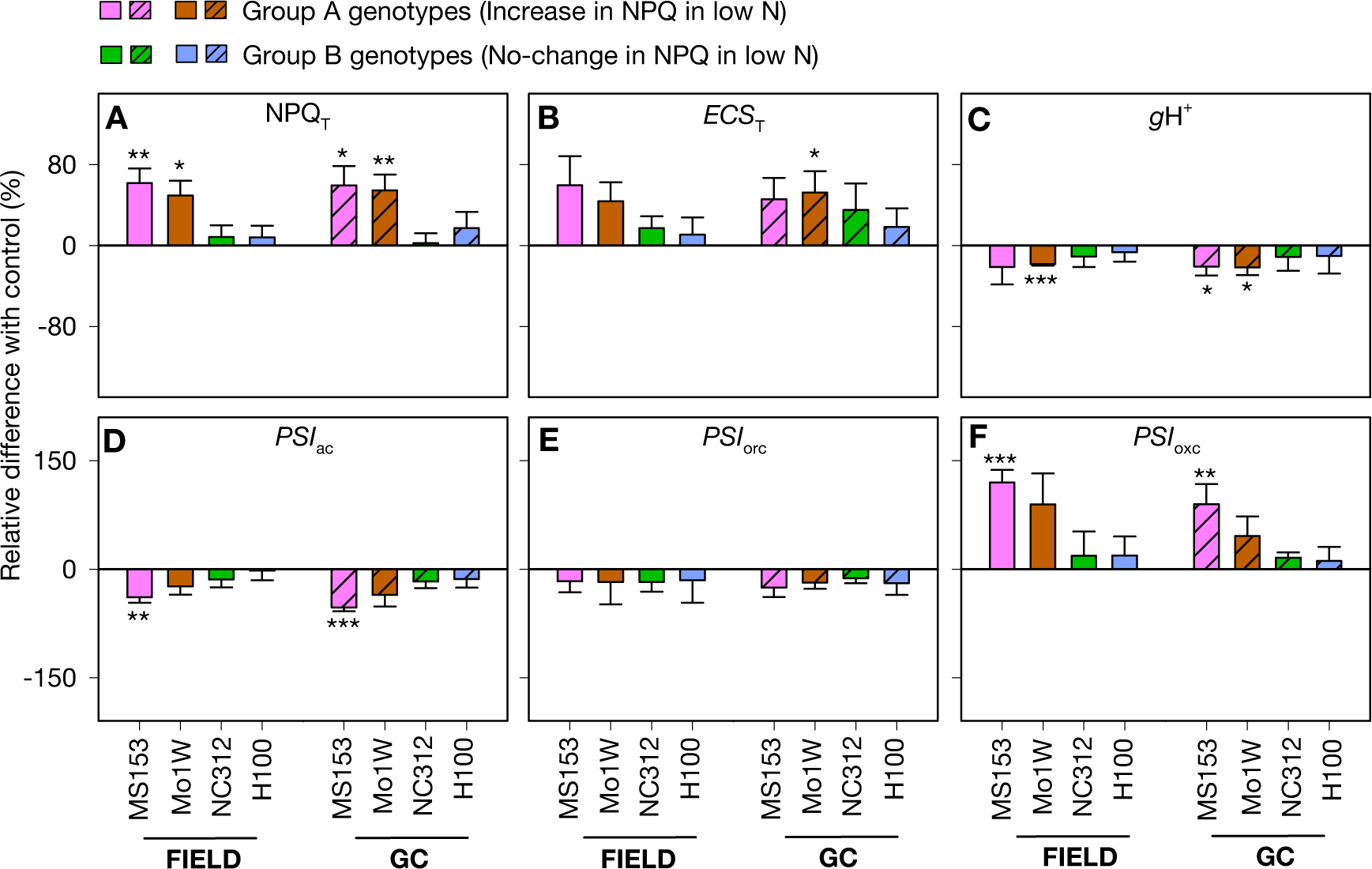
Photosynthesis-related traits in four maize genotypes grown on two nitrogen treatments in the growth chamber. **(A)** Theoretical NPQ (NPQ_T_). **(B)** Proton motive force delivered from the electrochromatic shift (ECS_T_). **(C)** proton conductivity related to ATPase activity (*g*H*^+^*). **(D)** PSI active centers (*PSI_ac_*). **(E)** PSI over-reduced centers (*PSI_orc_*), and **(F)** PSI oxidized centers (*PSI_oxc_*). Traits were measured in the late-vegetative stage in MS153, Mo1W (increase NPQ in low N; group A genotypes), and NC312 and H100 (no change in NPQ in low N; group B genotypes). Numbers for low N are shown as a relative difference (%) to control. Data represent mean ± SEM (n = 4 plots in 2020 field trail and n = 4 plants in growth chamber (GC)). Asterisks indicate significant differences with control (paired t-test; * *\*P* ≤ 0.05, *\*\*P* ≤ 0.01, \*\*\**P* ≤ 0.001 are given in Dataset 12.

**Fig S13.**
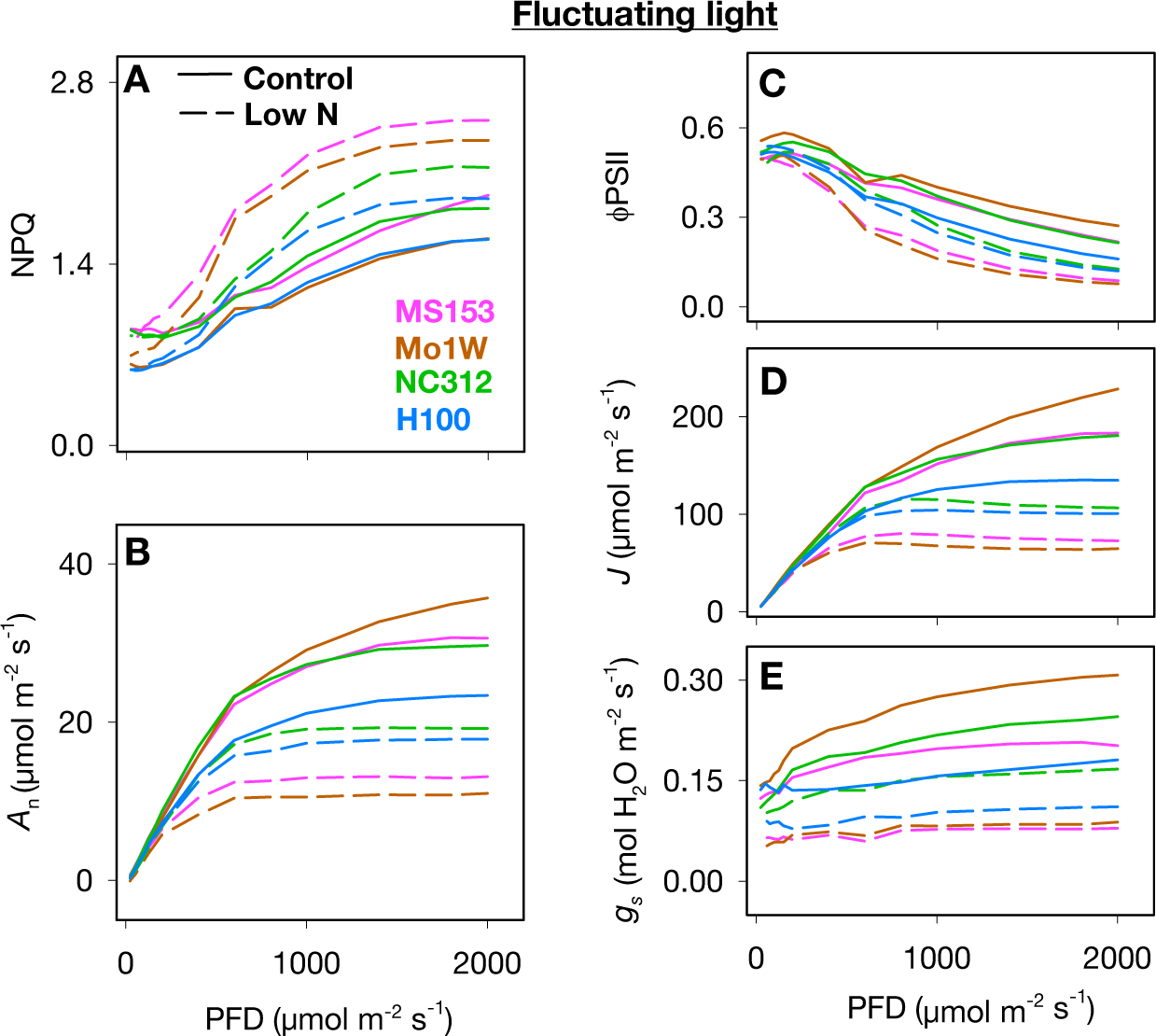
Response of photosynthesis-related parameters to fluctuating light conditions in four maize (*Zea mays* L.) genotypes under two nitrogen treatments performed in the growth chamber. **(A)** NPQ, **(B)** Photosystem II operating efficiency (ΦPSII), **(C)** Stomatal conductance (*g*_s_) **D)** Net CO_2_ fixation rate (*A*_n_), and **(E)** Linear electron transport rate (*J*), as a function of incident photon flux density (PFD). Traits were measured in the late-vegetative stage in MS153, Mo1W (increase in NPQ in low N; group A genotypes), and NC312 and H100 (no change in NPQ in low N; group B genotypes) in control (solid line) and low N (dashed line) conditions. Data represents means (n = 4 plants). Data used to produce this figure are given in Dataset 6.

**Fig S14.**
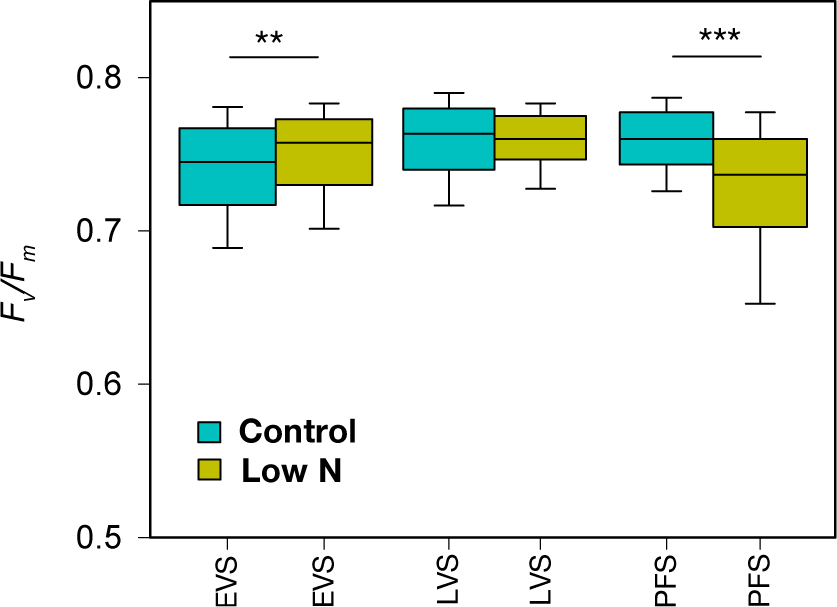
Maximum PSII quantum yield (*Fv/Fm*) at three developmental stages for maize genotypes grown in the field under control and low nitrogen conditions. *F_v_/F_m_* delivered as a first point of the assay in dark for 225 genotypes at early-vegetative (EV), late-vegetative (LV), and post-flowering (PF) stages measured in 2019 under control and low N conditions. The central line within each box plot represents the median; box edges indicate the upper and lower quantiles; upper and lower whiskers show minimum and maximum at the 1.5 × interquartile range (n = from 224 to 225 genotypes). Asterisks indicate significant differences between control and low N inside the developmental stage (paired t-test; \*\**P ≤ 0.01; ***P ≤* 0.001). Data used to produce this figure are given in Dataset 1.

**Table S1.** List of 225 genotypes in 2019 and a subset of 22 genotypes in 2020 from the maize (*Zea mays* L.) association used in the present study. As a result of its large size an excel (.xlsx) file for Table S1 is available online in the Supporting Information section.

**Table S2.** Schematic representations of the field experimental design in 2019. As a result of its large size an excel (.xlsx) file for Table S2 is available online in the Supporting Information section.

**Table S3.** Schematic representations of the field experimental design in 2020. As a result of its large size an excel (.xlsx) file for Table S3 is available online in the Supporting Information section.

**Table S4.** Derivative traits of non-photochemical quenching (NPQ) kinetics with their description used in the present study.

| Kinetics parameter | Description | Calculation function | Kinetics attribute |
| --- | --- | --- | --- |
| $NPQ_{slope_{light}}$ | Initial slope of NPQ in light | Hyperbola | Rate |
| $NPQ_{rate\_constant_{light}}$ | Rate in which NPQ reaches 63.2% of final steady state value ($\text{min}^{-1}$) during light | Exponential | |
| $NPQ_{asymptote_{light}}$ | Asymptote of NPQ during light | Exponential | Steady state |
| $NPQ_{max}$ | Maximum NPQ during light | The highest value of NPQ reached during 10 min illumination | |
| $NPQ_{start_{dark}}$ | NPQ at the beginning of the dark | Hyperbola | N/A |
| $NPQ_{rate\_constant_{dark}}$ | Rate in which NPQ reaches 63.2% of final steady state value ($\text{min}^{-1}$) during dark | Exponential | Rate |
| $NPQ_{amplitude_{darkE}}$ | Amplitude of NPQ during dark | Exponential | Range |
| $NPQ_{residual_{dark}}$ | Residual of NPQ in end of dark | Not relaxed NPQ in the end of the assay calculated from exponential function | Steady state |
| $F_v/F_m$ | Maximum PSII quantum yield | Measured before turning on the light (1<sup>st</sup> point of assay) | N/A |

**Supplemental Method 1 Hoagland Medium modification for N stress treatment**

In the preliminary experiment, four nitrogen treatments: control (10 mM NO_3-_), moderate (7.5mM NO_3-_), low (5 mM NO_3-_) and insufficient N (2.5 mM) were tested on B73 maize genotype. Control, moderate, low, and insufficient N treated plants were receiving 210.2 mg (5 mM KNO_3_ and 5 mM Ca(NO_3_)_2_.4H_2_O), 140.0 mg (5 mM KNO_3_ and 2.5 mM Ca(NO_3_)_2_.4H_2_O), 105.1 mg (2.5 mM KNO_3_ and 2.5 mM Ca(NO_3_)_2_.4H_2_O), and 35.0 mg (2.5 mM KNO_3_ only) nitrogen per liter (L) Hoagland media every day, respectively. Hoagland media was modified with different nitrogen treatments by replacing KNO_3_ with K_2_SO_4_ and Ca(NO_3_)_2_.4H_2_O with CaCl_2_ salts. The working Hoagland solution (1L) contained 2.5 to 5 mM KNO_3_, 0 to 5 mM Ca(NO_3_)_2_.4H_2_O, 2.5 mM K_2_SO_4_, 1 mM KH_2_PO_4_ (pH 6.0), 2 mM MgSO_4_.7H2O, 50 µM Fe-EDTA, 0.046 mM H_3_BO_3_, 0.009 mM MnCl_2_, 7.5×10^-4^ mM ZnSO_4_.7H_2_O, 3.2×10^-4^ mM CuSO_4_, and 1.1 × 10^-4^ mM H_2_MoO_4_.

**Dataset 1** NPQ kinetics data collected for 225 maize genotypes under two nitrogen treatments in the field of 2019.

**Dataset 2** NPQ kinetics data collected for a subset of 22 maize genotypes under two nitrogen treatments in the field of 2020.

**Dataset 3** Diurnal data of NPQ_T_ and other fluorescence related parameters collected for a subset of 22 maize genotypes under control and low nitrogen in the field of 2020.

**Dataset 4** Pigments related data collected for 10 maize genotypes in the field 2020 and 4 genotypes in the growth chamber.

**Dataset 5** NPQ kinetics data collected for 4 maize genotypes under control and low nitrogen in the growth chamber.

**Dataset 6** Gas exchange data from steady state and fluctuating A/Q response collected for 4 maize genotypes under control and low nitrogen in the growth chamber.

**Dataset 7** Gas exchange data from A/Ci response collected for 4 maize genotypes under control and low N in the growth chamber.

**Dataset 8** Data related to morphological parameters collected for 4 maize genotypes under control and low nitrogen in the growth chamber.

**Dataset 9** Data related to absorbance, fluorescence, and biomass parameters collected for B73 maize genotype under different nitrogen treatments in the growth chamber.

**Dataset 10** Data related to NPQ_T_ and other fluorescence parameters (ϕ_NPQ,_ ϕ_PSII,_ and ϕ_NO_) parameters collected for two groups (Group A and B) of maize genotypes under control and low nitrogen in field 2020.

**Dataset 11** NPQ kinetics data collected for 4 maize genotypes at different light intensities under two nitrogen treatments in the growth chamber.

**Dataset 12** Data related to absorbance and fluorescence parameters collected for 4 maize genotypes under two nitrogen treatments in the field and growth chamber.

## Author contributions

KG and JCS conceived the project. SS, MG and KG collected the data from the field trials. SS collected the data from growth chamber trials. SS analyzed the data. MG conducted additional analyses. SS and KG designed figures. JCS designed an additional figure. SS, KG and JCS drafted the manuscript. All authors read, edited, and approved the final manuscript.

## Acknowledgements and Funding

We would like to acknowledge Brandi Sigmon and Christine Smith for designing and executing the field experiments in 2019 and 2020. The authors thank Annie Nelson, Himani Patel and Marie Barnald for assistance in the collection of leaf disks from the field and imaging them in the lab. This research was supported by the National Science Foundation under award no. OIA-1557417 for the Center for Root and Rhizobiome Innovation (CRRI). KG was supported by the National Science Foundation supporting Nebraska Established Program to Stimulate Competitive Research program under FIRST Award, Layman Fund held at the University of Nebraska Foundation and National Science Foundation CAREER grant no. 2142993.

## Data availability

All data supporting the findings of this study are available online in the Supporting Information section of datasets (.xlsx).

